# TOR and heat shock response pathways regulate peroxisome biogenesis during proteotoxic stress

**DOI:** 10.1101/2024.12.31.630809

**Authors:** Nandini Shukla, Maxwell L. Neal, Jean-Claude Farré, Fred D. Mast, Linh Truong, Theresa Simon, Leslie R. Miller, John D. Aitchison, Suresh Subramani

## Abstract

Peroxisomes are versatile organelles mediating energy homeostasis and redox balance. While peroxisome dysfunction is linked to numerous diseases, the molecular mechanisms and signaling pathways regulating peroxisomes during cellular stress remain elusive. Using yeast, we show that perturbations disrupting protein homeostasis including loss of ER or cytosolic chaperone function, impairments in ER protein translocation, blocking ER N-glycosylation, or reductive stress, cause peroxisome proliferation. This proliferation is driven by increased *de novo* biogenesis from the ER as well as increased fission of pre-existing peroxisomes, rather than impaired pexophagy. Notably, peroxisome biogenesis is essential for cellular recovery from proteotoxic stress. Through comprehensive testing of major signaling pathways, we determine this response to be mediated by activation of the heat shock response and inhibition of Target of Rapamycin (TOR) signaling. Finally, the effects of proteotoxic stress and TOR inhibition on peroxisomes are also captured in human fibroblasts. Overall, our findings reveal a critical and conserved role of peroxisomes in cellular response to proteotoxic stress.

## INTRODUCTION

Peroxisomes are essential, evolutionarily-conserved organelles that play key roles in energy metabolism via fatty-acid oxidation and redox homeostasis ^1^. Their biogenesis is controlled by *PEX* genes, many of which are conserved across species, and their turnover occurs by selective autophagy, called pexophagy ^2-4^. Peroxisome biogenesis occurs either by growth and division of pre-existing peroxisomes, or by *de novo* biogenesis, which involves the budding and subsequent fusion of pre-peroxisomal vesicles from the endoplasmic reticulum (ER) ^2,5,6^. The balance between peroxisome biogenesis and turnover maintains the homeostasis of this dynamic organelle ^7-9^. In most organisms, peroxisome homeostasis is sensitive to nutritional cues. This is exemplified in single-celled yeasts by the induction of peroxisomes in cells grown in oleate because fatty acid oxidation is exclusively peroxisomal ^10,11^. In rodents, a diverse class of compounds called peroxisome proliferators induce adaptations consisting of hepatocellular hypertrophy and hyperplasia, and transcriptional induction of fatty-acid metabolizing enzymes coincident with peroxisome proliferation ^12^; however, the regulation of human peroxisomes remains unclear ^13,14^. Not surprisingly, significant efforts have focused on the intracellular signaling pathways responding to nutritional cues resulting in peroxisome induction ^10,15-18^.

However, peroxisomes also respond to environmental stresses, and proteins are known to traffic into the organelle in response to stress ^19^, but the studies investigating the signaling components and mechanisms involved are limited in both number and scope ^20-22^. The importance of peroxisomes arises not only from their metabolic specialization, but also from their roles in immunometabolism, development, aging and human disease ^23-28^, most notably in the context of human peroxisome biogenesis disorders (PBDs). This is the driving imperative to understand peroxisome biogenesis and its modulation, particularly its role in the adaptation to varying forms of abiotic environmental conditions, such as heat ^21,29-31^, light ^32^, pH ^33^, salt ^22^, redox ^34,35^ and organelle stress ^20^, which necessitate a coordinated cellular response, often requiring contributions from several sub-cellular compartments and membrane contact sites^2,36,37.^

In this study, we demonstrate that proteotoxic ER stress induced by tunicamycin treatment causes peroxisome proliferation in *Saccharomyces cerevisiae, Komagataella phaffii*, and primary human fibroblasts. By manipulating the many arms of the ER stress response either genetically or pharmacologically, we discern that proteotoxic ER stress increases peroxisome number partially by activating the heat shock response (HSR) in the cytosol and inhibiting TOR1. Mechanistically, this is executed through increased peroxisome production via mainly the *de novo* mode along with fission of pre-existing peroxisomes, instead of impaired pexophagy.

Importantly, peroxisome biogenesis is necessary for cell survival in response to tunicamycin treatment. Our study thus sheds light on the role of *de novo* peroxisome biogenesis in cellular adaptation to ER stress, with emphasis on evolutionary conservation of this adaptation, the role of multiple sub-cellular compartments in coordinating the response, as well as the types of stress and signaling pathways that activate this proliferation response. Overall, this study paves the path for a better understanding of peroxisome biogenesis in response to environmental insults, while also presenting potential applications in the management of PBDs.

## RESULTS

### Inactivation of Kar2/BiP triggers UPR and causes peroxisome proliferation

While analyzing essential *S. cerevisiae* mutants for potential peroxisomal phenotypes, such as altered number using the fluorescent peroxisomal matrix marker, GFP-ePTS1, we observed that the temperature-sensitive mutant, *kar2-159*, exhibited a strikingly elevated number of peroxisomes compared to WT. A more quantitative analyses of peroxisome number from 3D images performed using our recently developed software, perox-per-cell ^38^, revealed that both, the WT and the *kar2-159* mutant cells, showed a wide distribution in peroxisome number at permissive (25°C) and restrictive (37°C) temperatures. However, *kar2-159* cells consistently displayed significantly more peroxisomes at 37°C, than at 25°C. In comparison, peroxisome proliferation was significantly, but only mildly, induced in WT cells in response to the temperature shift (Figure 1a-b).

**Figure 1:**
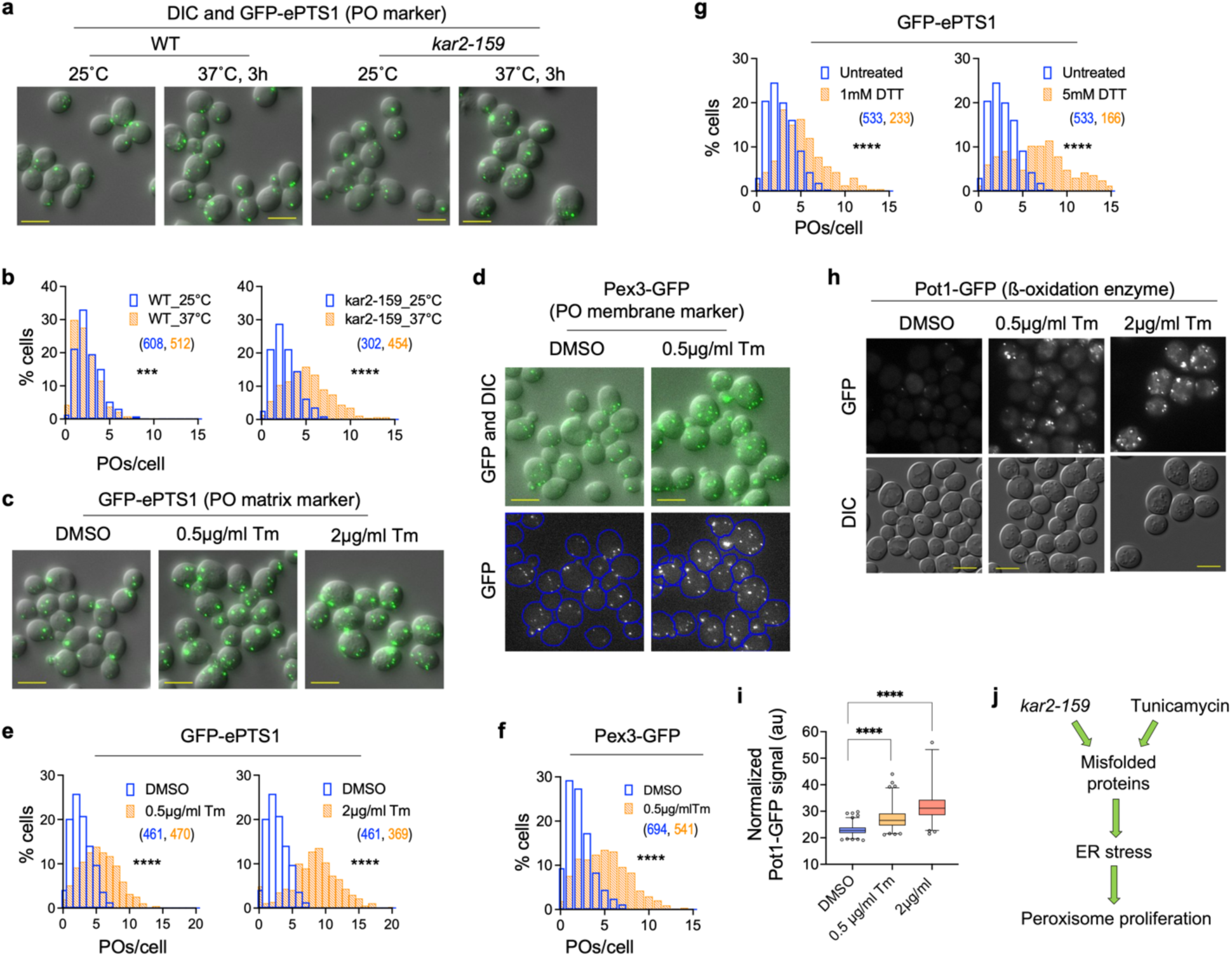
Protein misfolding stress induces peroxisome proliferation. a) Z-projection images showing peroxisomes marked by GFP-ePTS1 in WT and *kar2-159* cells grown at 25°C and for 3h after transfer to 37°C. b) Histograms showing population distribution of the number of peroxisomes per cell (POs/cell) in WT and *kar2-159* mutant cells at 25°C and after 3h of growth at 37°C. The *kar2-159* mutant cells have a higher number of peroxisomes at the restrictive temperature (37°C) compared to WT. c-f) Z-projection images (c-d) and histograms (e-f) showing the number of peroxisomes at 5h after treatment with DMSO or tunicamycin (Tm, 0.5µg/mL or 2.0µg/mL as shown). Peroxisomes were visualized using either a peroxisomal matrix marker, GFP-ePTS1 (c, e), or a peroxisomal membrane protein, Pex3-GFP (d, f). Tunicamycin treatment increases the number of peroxisomes per cell as measured by both markers. g) Histograms showing increased peroxisome abundance at 5h after treatment with 1mM or 5mM DTT. h-i) Single Z-slice images (h) and quantification (i) showing that Pot1-GFP (peroxisomal thiolase) levels per cell increase after treatment with tunicamycin (number of cells, i.e., N_cells_: DMSO: 510, 0.5µg/ml Tm: 454, 2µg/ml Tm: 340). j) Schematic depicting that cells experiencing ER stress, which could be caused either by loss of Kar2 function or treatment with tunicamycin, manifest peroxisome proliferation phenotypes, such as increased organelle number and increased level of peroxisomal enzymes that are typically repressed during growth in glucose. In this and subsequent figures, the number of cells for each histogram is indicated in parentheses and depicted in matching font color as the histograms, and statistical analyses of population distributions before and after treatments (e.g. tunicamycin/other drugs vs DMSO or 25°C vs 37°C), or comparing peroxisome number in mutants vs WT, were performed using the Mann-Whitney test (*, p<0.05; **, p<0.01; ***, p<0.001; ****, p<0.0001). Scale bar: 5µm.

Kar2, the yeast homolog of mammalian BiP, belongs to the Hsp70 family of chaperone proteins residing in the ER lumen ^39,40^. During protein misfolding stress in the ER lumen, Kar2 binds to unfolded proteins thereby dissociating from its binding partner, Ire1. This causes the activation of the unfolded protein response (UPR), a transcriptional response mediated by Hac1, which binds to the UPR elements (UPREs) of gene promoters to activate expression of genes for chaperone function, ER expansion and ER associated degradation (ERAD). We verified that inactivation of Kar2 induces the UPR (quantified by UPRE-GFP) ^41,42^ (Supplemental Figure 1a-b), as well as the cytosolic heat shock response (quantified by HSE-GFP) ^42,43^ (Supplemental Figure 1c-d), under our experimental conditions. To understand how loss of Kar2 function is connected to peroxisome homeostasis, we investigated the effects of UPR and heat shock response activation on peroxisome proliferation.

### Induction of proteotoxic, but not lipid bilayer, ER stress causes peroxisome proliferation and upregulates peroxisomal 3-ketoacyl-CoA thiolase

After verifying under our assay conditions, the induction of UPRE-GFP by tunicamycin (Supplemental Figure 1e-f) (Tm), a nucleoside antibiotic that interferes with N-glycosylation of proteins on the luminal side of the ER, we tested its effect on peroxisome number by counting Pex3-GFP and GFP-ePTS1 puncta. Pex3 is a peroxisomal membrane protein (PMP), whereas GFP-ePTS1 is imported into the peroxisome lumen. Tunicamycin significantly increased the number of peroxisomes as visualized by both markers (Figure 1 c-f) within 3-5h. ER stress induced by DTT treatment also increased peroxisome number (Figure 1g). Our findings align with a recent report, which suggested that peroxisome numbers (monitored indirectly by total Pex11 fluorescence) ^20^ increase as an adaptation to supply additional acetyl-CoA via peroxisomal ß-oxidation to support mitochondrial respiration during ER stress. To directly explore this idea, we examined whether ER stress induced peroxisomal β-oxidation enzymes, which are typically repressed during growth in glucose. We found that the endogenous levels of GFP-tagged, 3-ketoacyl-CoA thiolase (Pot1) ^18^, which catalyzes the final step in peroxisomal ß-oxidation, increased following tunicamycin treatment (Figure 1h-i). Collectively, our observations indicate that ER stress resulting from misfolded proteins, caused either by loss of Kar2 function or by tunicamycin-mediated protein glycosylation defects, results in cellular peroxisome proliferation (Figure 1j, Supplemental Figure 1).

To investigate if UPR induction causes peroxisome proliferation, we abrogated the UPR by either deleting Ire1 or Hac1. Notably, another transcription factor, Gcn4, is also upregulated by ER stress, can bind to UPREs, and is required for the induction of several UPR target genes, including *PEX* genes (*PEX11*, *PEX14*, *PEX21*) involved in peroxisome biogenesis and their regulators (*PXA2*, *PIP2*) ^44-46^. Tunicamycin treatment induced peroxisome proliferation in *ire1Δ*, *hac1Δ* and *gcn4Δ* cells, indicating that a functional UPR is not required for peroxisome proliferation during ER stress (Figure 2a-c). Lipid imbalance also causes ER stress ^47^, therefore we induced lipid bilayer stress (LBS) by growing cells in media lacking inositol to assess its effects on peroxisomes. While growth of yeast in media lacking inositol induced UPR (Supplemental Figure 2a) and had a mild effect on HSE-GFP level (Supplemental Figure 2b), there was no significant increase in the number of peroxisomes (Figure 2d) and only a moderate increase of Pot1-GFP levels (Figure 2e). In response to ER stress, cells also activate the ER stress surveillance (ERSU) pathway which prevents the inheritance of damaged ER into daughter cells ^48^. Activation of the ERSU pathway, using phytosphingosine, an early biosynthetic sphingolipid, did not increase the number of peroxisomes (Figure 2f), Pot1-GFP levels (Figure 2g), UPR induction (Supplemental Figure 2c) or HSE-GFP levels, to the extent seen following tunicamycin treatment (Supplemental Figure 1g-h, 2d). These observations suggest that proteotoxic ER stress, but not lipid bilayer stress, causes peroxisome proliferation in a UPR- and ERSU-independent manner (Figure 2h).

**Figure 2:**
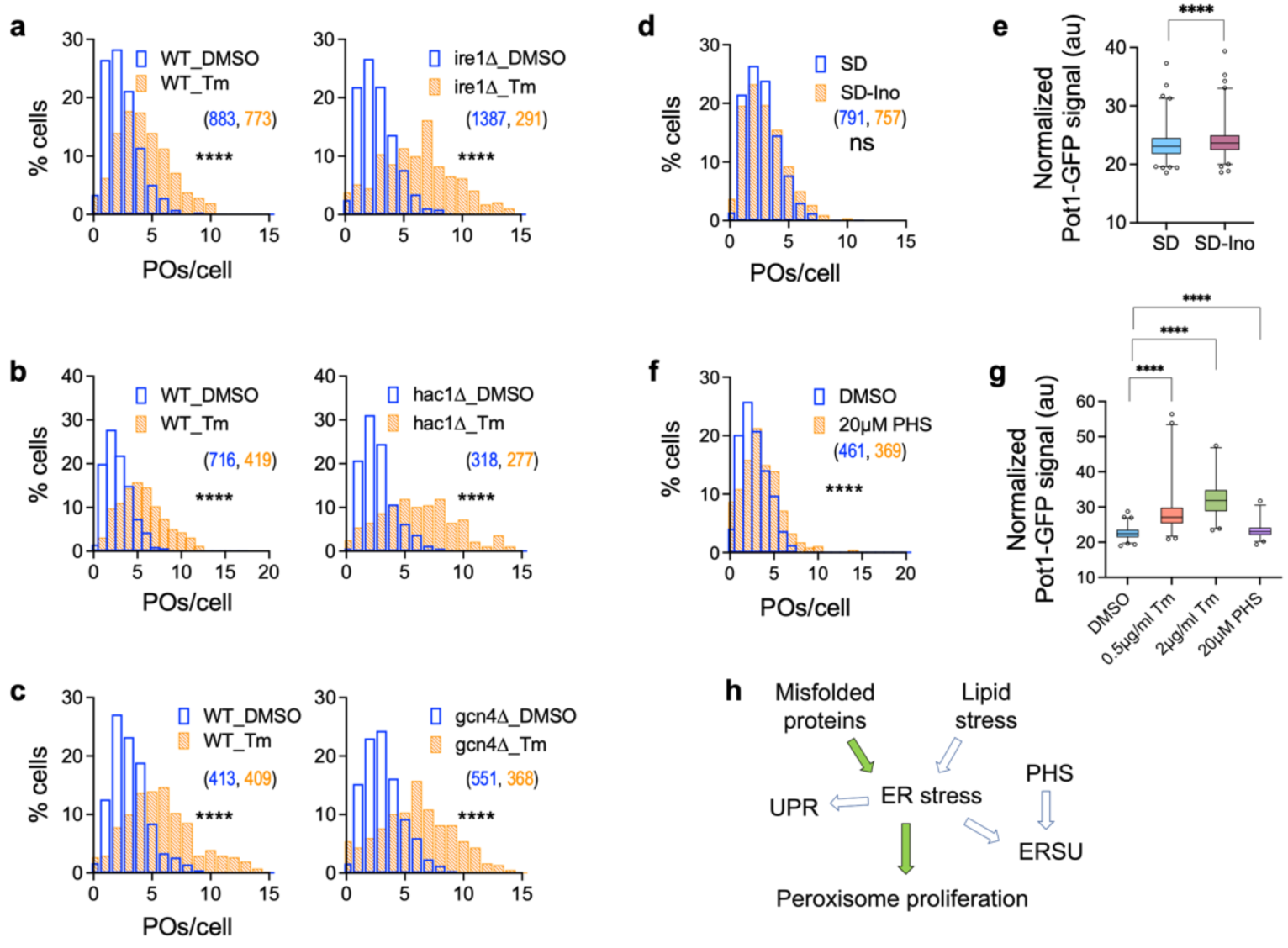
ER stress-induced peroxisome proliferation occurs independent of the UPR pathway. a-c) Histograms showing that tunicamycin-induced peroxisome proliferation in the UPR pathway mutants, such as *ire1*Δ, *hac1*Δ and *gcn4*Δ, is not reduced relative to WT cells. d-e) Histogram of peroxisomes per cell (d) and box plot showing normalized Pot1-GFP levels (e) demonstrating that in WT cells grown in the absence of inositol, neither feature increases as dramatically as after tunicamycin treatment (see Figure 1c,e,h,i), during growth in the absence of inositol (N_cells_: SD: 523, SD-Ino: 405). f-g) Histogram of number of peroxisomes per cell (f) and box plot for normalized Pot1-GFP levels (g) indicating that neither show an increase comparable to tunicamycin treatment, after treatment with Phytosphingosine (PHS) (N_cells_: DMSO: 327, 0.5µg/ml Tm: 264, 2µg/ml Tm: 227, 20µM PHS: 228). h) Flow chart showing the potential routes for transmission of signals to induce peroxisome proliferation during stress. The green arrows show the routes which we experimentally verified to be used for transmission, whereas the clear arrows show the path which we verified to either not be required, viz. UPR, or not be sufficient, viz. Lipid stress and PHS treatment for inducing peroxisome proliferation.

### Misfolded protein stress causes peroxisome proliferation partially via heat shock response activation

Kar2 also functions in the passage of proteins across the Sec61-Sec63 translocon in the ER membrane ^49^. By binding to the translocation substrate polypeptide, Kar2 acts as a molecular ratchet to prevent its back-translocation, thereby facilitating its unidirectional movement only towards the ER lumen ^50,51^. We investigated if peroxisome proliferation observed in *kar2-159* cells is due to impaired ER-protein translocation and subsequent heat shock response activation (Figure 3a). We blocked ER protein targeting, without directly interfering with Kar2 or the Sec61-Sec63 translocon, by impairing the GET pathway, which mediates the insertion of Tail-anchored (TA) proteins into the ER ^52^. We observed that peroxisome number increased in the absence of Get3 (Figure 3b-c) as also noted in a recent study on organelle tethering ^53^. Absence of Get3 also increased HSE-GFP levels at all temperatures examined between 25-37°C (Figure 3d, Supplemental Figure 3a), indicating the activation of heat shock response by mistargeted TA proteins.

**Figure 3:**
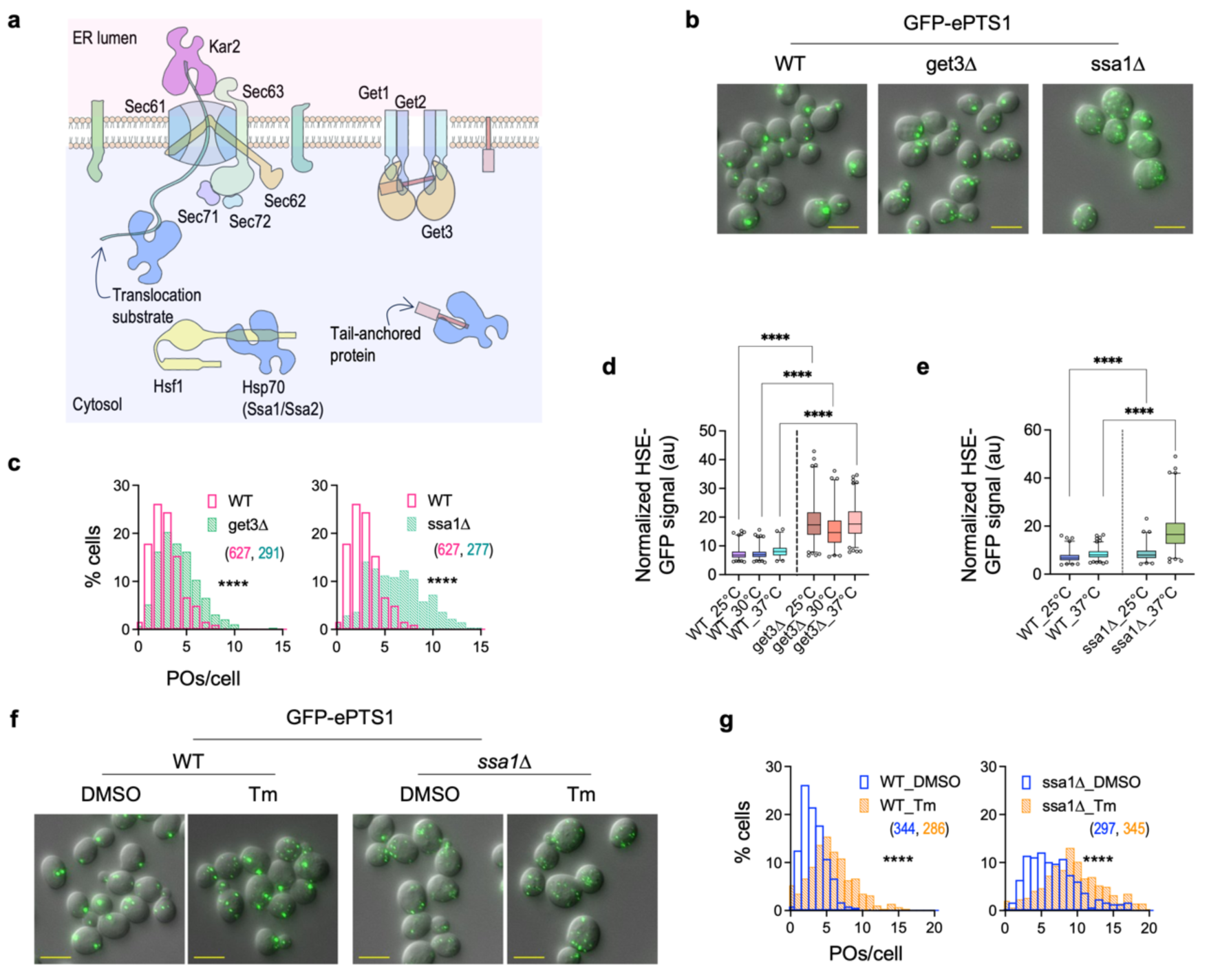
Peroxisome proliferation seen during ER stress is partially driven via heat shock response activation. a) Cartoon showing molecular players involved in the crosstalk between protein homeostasis in the ER and cytosol. Proteins destined for the ER lumen are translocated across the ER membrane via the Sec61 or Get pathways. Kar2 on the luminal side and Hsp70s including Ssa1 in the cytoplasm prevent the translocation substrates from misfolding. Hsp70s also keep tail-anchored proteins from misfolding in the cytoplasm before their transfer to Get3 for membrane insertion. Hsp70s additionally are bound to Hsf1 and keep it inactive; however, increased burden of unfolded proteins in the cytoplasm can titrate Hsp70 away from Hsf1, thereby activating the latter. Protein translocation defects in *kar2-159* at 37°C, or in *get3*Δ can potentially cause the buildup of ER proteins in the cytoplasm, thereby activating the heat shock response. b-c) Z-projection images (b) and histograms (c) showing that cells lacking Get3 or Ssa1 show increased peroxisome proliferation. d-e) Deletion of *GET3* (d), or of *SSA1* at 37°C (e) activates the heat shock response [(d) N_cells_ for WT: 25°C: 504, 30°C: 540, 37°C: 270; N_cells_ for *get3*Δ: 25°C: 579, 30°C: 316, 37°C: 537; (e) N_cells_ for WT: 25°C: 416, 37°C: 620; N_cells_ for *ssa1*Δ: 25°C: 344, 37°C: 491)]. f-g) Z-projection images (f) and histograms (g) showing that peroxisome proliferation seen in *ssa1*Δ is comparable to that induced by tunicamycin. Tunicamycin also further increases the number of peroxisomes in *ssa1Δ* cells.

Analogous to the UPR in the ER, the heat shock response pathway is activated in response to the presence of misfolded and aggregated proteins in the cytosol. Signaling through this pathway is primarily mediated by Hsf1, which binds to the Heat Shock Elements (HSEs) in gene promoters and activates transcription of chaperones and Ubiquitin-proteasome system (UPS)-mediated degradation ^54^. We induced the heat shock response pathway by deleting the *SSA1* gene, encoding the Hsp70 binding partner of Hsf1 ^30^. Typically, the Hsp70 chaperones Ssa1 and Ssa2 keep Hsf1 inactive; however, under stress, unfolded or misfolded proteins titrate the Hsp70s away from Hsf1, thereby allowing nuclear translocation of Hsf1 and transcriptional activation at the HSEs. The *ssa1*Δ cells had not only elevated HSE-GFP levels (Figure 3e, Supplemental Figure 3b), but also increased number of peroxisomes compared to WT (Figure 3b-c). While the median number of peroxisomes in *ssa1*Δ was 7-8 compared to 3 in WT, ∼70% of *ssa1*Δ cells had >5 peroxisomes, compared to ∼5% in WT. The absence of Ssa1 caused a comparable increase in the number of peroxisomes as seen after tunicamycin treatment (Figure 3f-g). Surprisingly, tunicamycin further increased the number of peroxisomes in *ssa1*Δ cells, with the median rising to 9-12 after tunicamycin treatment, compared to 6-7 in DMSO-treated cells (Figure 3f-g). This suggests that tunicamycin and Hsf1 activation may act synergistically in stimulating peroxisome proliferation.

We compared the effect of tunicamycin on *ssa1*Δ vs. WT cells using a generalized linear modeling (GLM) approach (see Materials and Methods). Briefly, peroxisome count data was fit to a hurdle model ^55^, using a GLM model formula containing terms that capture the effects of experimental batch, strain and treatment, as well as an interaction term that captures the strain-specific response to treatment. We then examined the value of the GLM coefficient on the interaction term to compare mutant vs. WT responses to tunicamycin. The statistical significance of these comparisons was estimated using likelihood ratio tests that compared GLM results using the full model formula to results from a reduced model where the interaction term is omitted. While simply comparing the median number of cellular peroxisome counts between strains in an experimental batch can provide a cursory glimpse into the relative effects of tunicamycin on those strains, our GLM approach allowed us to account for several important features of our data to increase the statistical power, resolution, and completeness of our comparisons. First, our experimental batches consisted of 1-4 mutant strains and a WT control. Using GLM with a model formula that includes a term for experimental batch effects allowed us to compare peroxisome counts between a mutant strain of interest and WT, using data pooled from all experimental batches, thereby maximizing statistical power. Second, some mutant strains had basal cellular peroxisome count distributions that were noticeably different from WT. Our GLM approach was used to ensure that these basal differences were accounted for when comparing mutant and WT tunicamycin responses. Third, whereas comparisons of median peroxisome counts have a limited resolution of 0.5, our GLM approach allowed us to quantify tunicamycin responses with a continuous variable. Our analysis revealed that the extent of peroxisome proliferation induced by tunicamycin in *ssa1*Δ cells was less than that in WT cells, suggesting that the loss of Ssa1 and the addition of tunicamycin promote peroxisome proliferation through partially overlapping mechanisms (Figure 4g).

**Figure 4:**
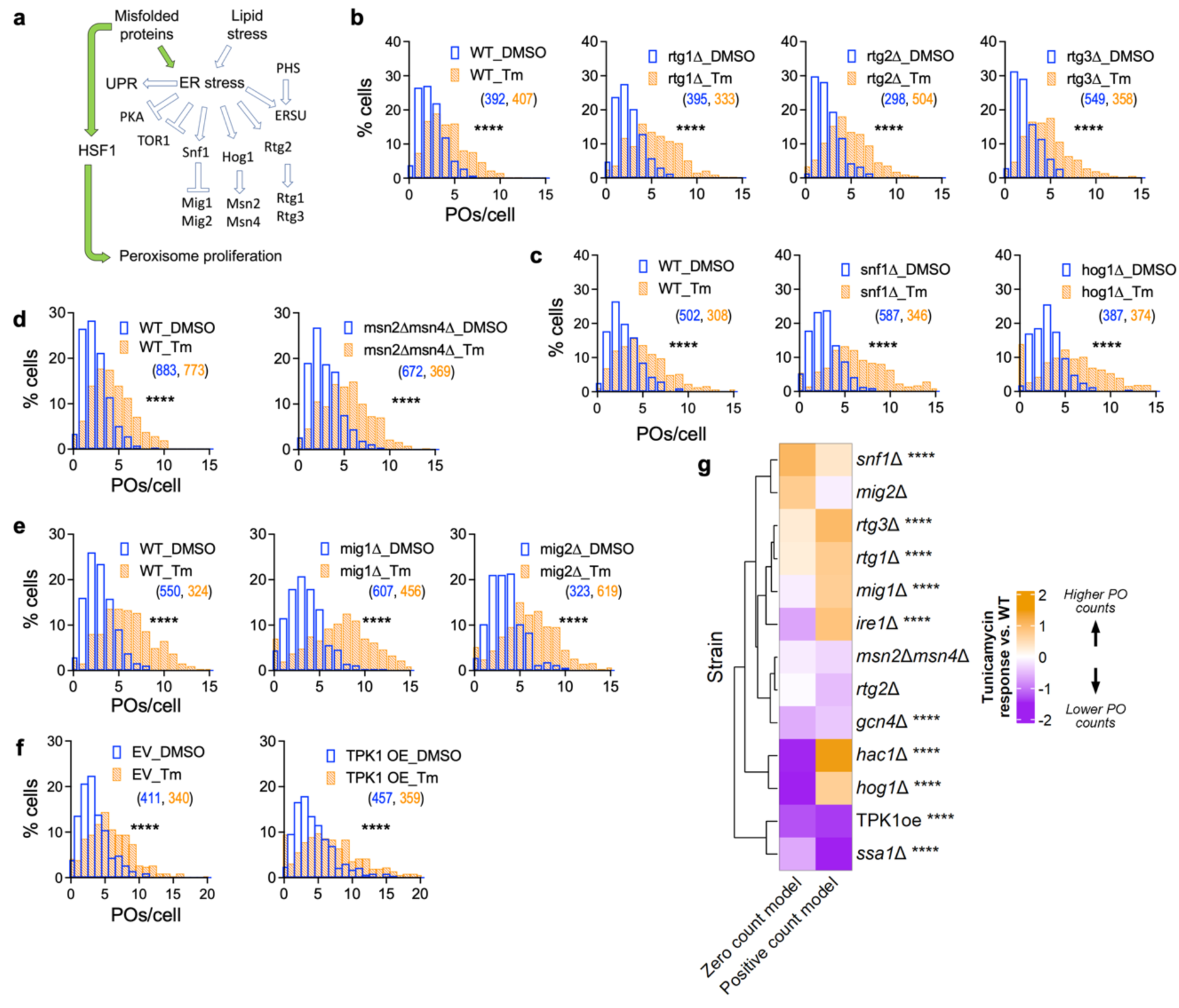
Peroxisome proliferation in response to misfolded protein stress occurs despite individual inactivation of Snf1, Hog1, and Rtg pathways. a) Flow chart showing a simple picture of the pathways that get activated or inhibited in response to ER stress without depicting the crosstalk among them. For detailed understanding see Figure 7. b-f) Histograms showing quantification of the number of peroxisomes per cell after treatment with tunicamycin or DMSO in mutants blocked in signaling through various pathways implicated in peroxisome biogenesis. Abrogating signaling through retrograde signaling (RTG) (b) Snf1-(c), and Hog1/Msn2/Msn4-(c,d) pathways, does not significantly prevent peroxisome proliferation in response to tunicamycin. Furthermore, the absence of transcriptional repressors, *mig1*Δ and *mig2*Δ, of peroxisome biogenesis genes (e) also does not mimic the effects of tunicamycin on peroxisome number. f) Constitutive activation of PKA by TPK1 overexpression (TPK1oe) using a multicopy plasmid does not fully block peroxisome proliferation in tunicamycin-treated cells. EV denotes empty vector control. g) Heat map of GLM analysis showing the mutants with reduced response in purple and those with increased response to tunicamycin in orange, compared to WT. The zero count and positive count models show the analyses of cells with zero and non-zero peroxisomes, respectively. *ssa1Δ* and TPK1oe strains are the only mutants that show significant and dramatically reduced peroxisome proliferation upon tunicamycin treatment in comparison to WT, whereas *gcn4Δ* showed significant, but relatively smaller, reduction.

### Peroxisome proliferation in response to ER stress is partially mediated by TOR1 inactivation

The adaptive response to ER stress occurs via activation of SNF1-, HOG1- and RTG-signaling pathways and inactivation of TOR1 and PKA ^45,46,56-61^; however, among these, the specific pathways that mediate the signaling response to trigger peroxisome proliferation in response to tunicamycin remain unclear (Figure 4a). We systematically abrogated each pathway and investigated the effect of tunicamycin on peroxisome proliferation in mutants compared to WT using our GLM analysis (Figure 4). Given published findings on the involvement of these pathways in yeast peroxisome proliferation and regulation of *PEX* gene expression, we were surprised to discover that blocking the retrograde signaling (RTG) (Figure 4b,g) ^22,62^, SNF1 ^10^ (Figure 4c,g), or HOG1 pathways ^17^ (Figure 4c,g) did not significantly impair peroxisome proliferation upon tunicamycin treatment. Similarly, the loss of both Msn2 and Msn4 (Figure 4d,g), which mediate the transcriptional response upon activation of the Hog1 pathway, as well as the heat shock response pathway ^17^, also did not prevent tunicamycin-induced peroxisome proliferation. Furthermore, we also found that cells lacking Mig1 or Mig2, peroxisome biogenesis repressors that are translocated from the nucleus by Snf1 activation ^63,64^, showed a comparable number of peroxisomes to WT, after both DMSO as well as tunicamycin treatments (Figure 4e,g). Consistent with our findings that peroxisome proliferation in ER stress is independent of Snf1, we observed that Snf1 was not activated after tunicamycin treatment under our assay conditions; however, Hog1 was activated after 4h (Supplemental Figure 4).

Response to ER stress also involves the inhibition of PKA ^61^ and TOR1 ^62^ (Supplemental Figure 4). We tested the role of PKA by using strains overexpressing TPK1 (TPK1oe) using a multicopy plasmid ^65,66^, which constitutively activates PKA, relative to a control strain carrying an empty vector (EV). Our GLM analysis indicated that the relative increase in peroxisome number in response to tunicamycin treatment was reduced in TPK1oe cells compared to that in cells carrying the empty vector (Figure 4f-g). This result suggests that PKA overexpression attenuates the tunicamycin-induced peroxisome proliferation.

Finally, we tested the effect of TOR1 inhibition on peroxisomes. After rapamycin treatment, while, we observed a mild increase in the number of peroxisomes in *S. cerevisiae* (Figure 5a-b), the effect in the methylotrophic yeast, *K. phaffii*, was significantly greater, exceeding the increase seen with tunicamycin addition, as indicated by a longer right tail of the distribution in rapamycin-treated, compared to tunicamycin-treated, cells (Figure 5c-e). Notably, the effects of tunicamycin and rapamycin in *K. phaffii* required twenty-fold higher (10µg/ml) drug concentration relative to *S. cerevisiae*, presumably due to robust efflux systems or reduced uptake efficiency in the former. However, we confirmed that the relative lower efficacy of rapamycin in *S. cerevisiae* was not due to sub-optimal drug concentration because increasing the rapamycin concentration (5µg/ml) did not increase peroxisome proliferation (Figure 5a-b).

**Figure 5:**
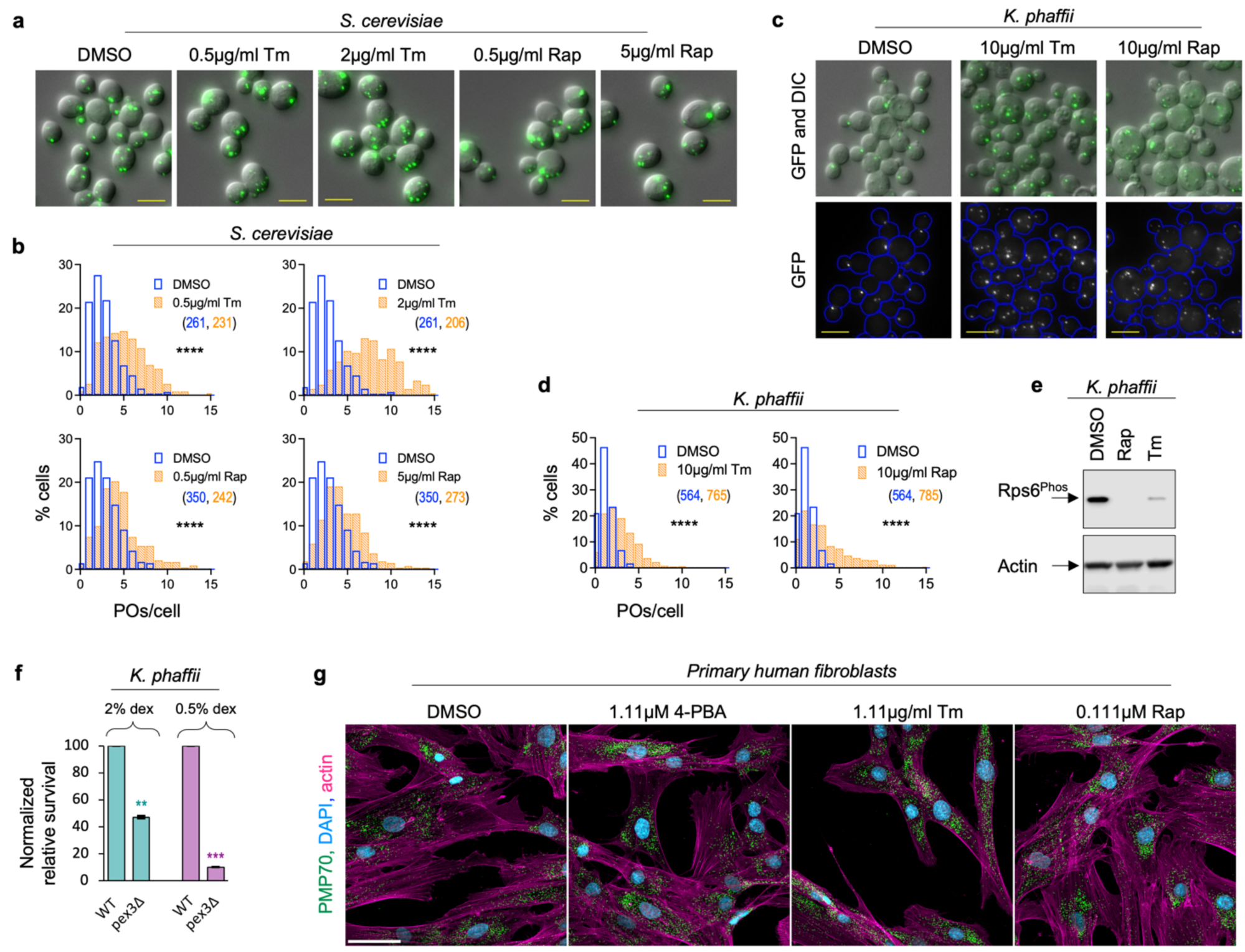
Misfolded protein stress causes peroxisome proliferation in a methylotrophic yeast via inactivation of Tor1. a-b) Z-projection images (a) and histograms (b) showing Tor1 inactivation by rapamycin treatment only mildly increases the number of peroxisomes in *S. cerevisiae*, compared to tunicamycin treatment, even with higher levels of rapamycin (5µg/ml). c-d) Z-projection images (c) and histograms (d) showing that tunicamycin as well as rapamycin causes peroxisome proliferation in the methylotrophic yeast, *K. phaffii*. e) Western blots showing that Rps6 phosphorylation decreases in cells treated with rapamycin or tunicamycin for 4h in comparison to DMSO in *K. phaffii*, indicating Tor1 inactivation after the two drug treatments. f) Quantification of survival of WT and *pex3*Δ *K. phaffii* cells after overnight treatment with 0.5µg/ml tunicamycin followed by treatment with 20µg/ml tunicamycin for 6h. Control treatments were performed with DMSO, and colony counts obtained for tunicamycin treatments were divided by those for DMSO treatments to obtain relative survival. Relative survival was normalized with WT to obtain the normalized relative survival for *pex3*Δ. Values indicate average from two experiments and error bars indicate standard deviation. Asterisks indicate statistical comparison of *pex3Δ* vs WT. g) Z-projection images showing that 1.11µg/ml tunicamycin, 0.111µM rapamycin and 1.11µM 4-PBA treatments increase peroxisome number in primary human fibroblasts. Nuclei (blue), peroxisomes (green) and actin (magenta) visualized using DAPI, PMP70-AF488 and Phalloidin-AF750 staining respectively. Scale bar: 50µm.

We tested the significance of peroxisomes in adaptation to ER stress by measuring the survival of WT and *pex3*Δ cells on rich media containing 2% or 0.5% dextrose after overnight (∼14h) treatment with a low dose of 0.5µg/ml tunicamycin followed by a high dose of 20µg/ml tunicamycin for 6h (Figure 5f). Cells lacking Pex3, which is essential for peroxisome biogenesis ^2^, consistently showed <50% survival in the tunicamycin-viability assay, which was reduced to <10% under reduced glucose availability compared to WT cells, underscoring the significance of peroxisome biogenesis in the adaptation.

To investigate the conservation of the peroxisome response to ER stress, we treated primary human fibroblasts with tunicamycin, rapamycin, or 4-phenyl butyric acid (4-PBA), a known inducer of peroxisomes, for 12h. Cells were fixed, stained for PMP70 (also known as ABCD3), a peroxisomal ATP-binding cassette (ABC) transporter that facilitates the transport of acyl-CoA esters across the peroxisomal membrane, and imaged (Figure 5g). Compared to DMSO-treated controls, 4-PBA increased the number of peroxisomes per cell. Likewise, tunicamycin treatment resulted in a dramatic increase in peroxisome numbers, with peroxisomes densely distributed throughout the cytoplasm. Rapamycin treatment produced mixed results, with many cells displaying elevated peroxisome numbers while others resembled the DMSO-treated controls. These findings demonstrate that the peroxisome proliferation response to ER stress is conserved in human cells. Overall, our findings indicate that among the many signaling pathways that are activated by ER stress, activation of the heat shock response pathway and inactivation of TOR1 and PKA are the primary mechanisms causing peroxisome proliferation.

### ER-stress driven peroxisome proliferation occurs by *de novo* biogenesis as well as growth and division

We systematically analyzed the contributions of peroxisome biogenesis, via growth and division and *de novo* pathways, and peroxisome loss by pexophagy, to ER-stress mediated peroxisome proliferation (Figure 6a). We reasoned that impairments in pexophagy would contribute to increase peroxisome number if: i) loss of the pexophagy receptor, Atg36 ^67^, phenocopies the effects of tunicamycin treatment on the number of peroxisomes without any drug, or ii) ER stress fails to further increase the number of peroxisomes in *atg36*Δ cells. While the absence of Atg36 did not increase basal peroxisome number, *atg36*Δ cells exhibited higher number of peroxisomes than WT after tunicamycin treatment (Figure 6b,d). These findings suggest that some peroxisomes are likely turned over during ER stress.

**Figure 6:**
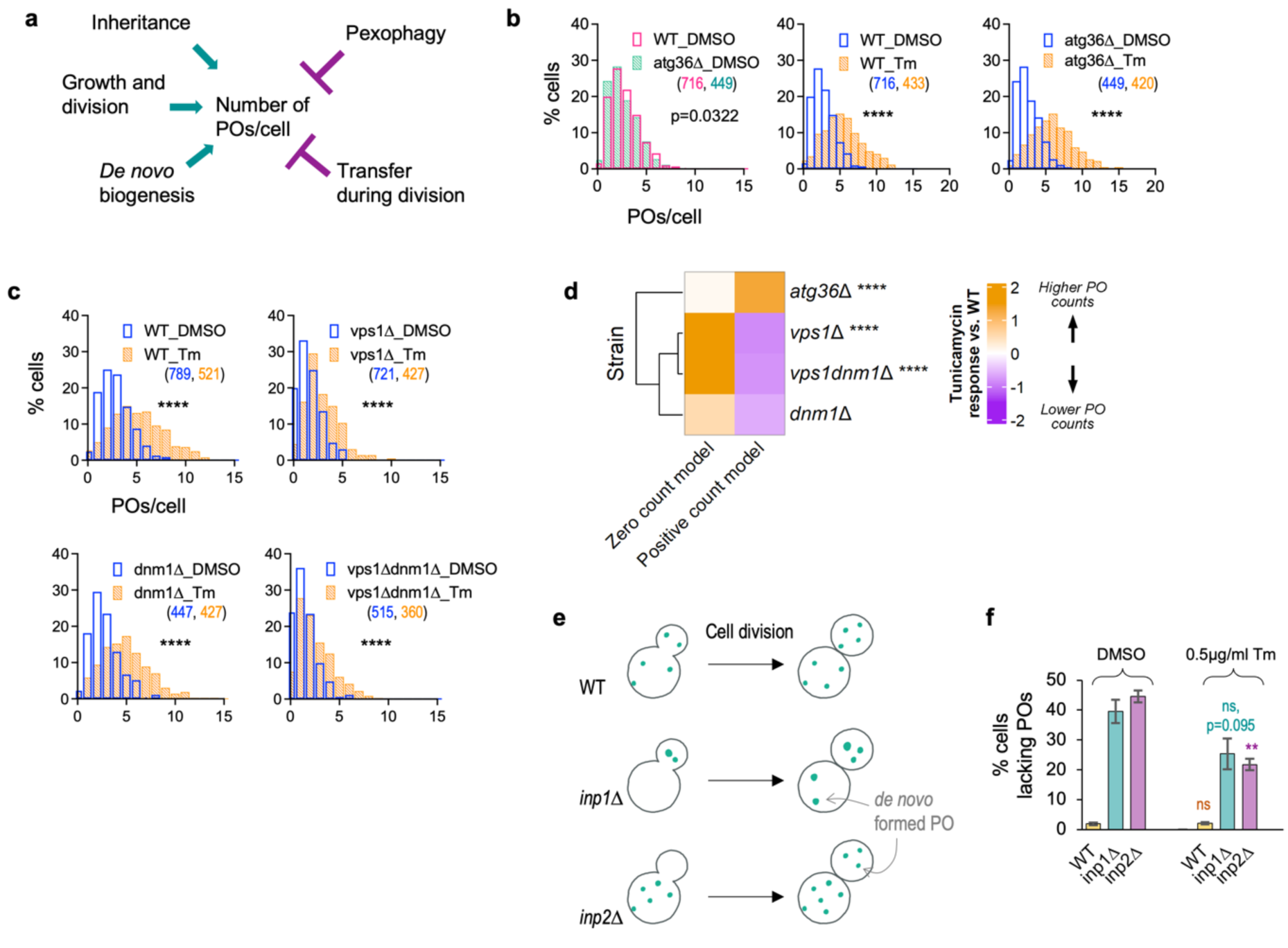
Increased peroxisome number during ER stress occurs by growth and division as well as *de novo* biogenesis. a) Cartoon showing that the peroxisome number in cells is regulated by a balance between pathways that increase peroxisomes, such as growth and division, *de novo* biogenesis from the ER and acquisition of peroxisomes due to inheritance, and those that reduce peroxisomes, such as pexophagy or transmission to daughter cells during cell division. b) Histograms showing that blocking pexophagy by deleting *ATG36* does not mimic tunicamycin-driven peroxisome proliferation. Deletion of Atg36 also does not impair peroxisome proliferation after tunicamycin treatment. c) Histograms showing cells can increase peroxisome number during ER stress in *vps1*Δ, *dnm1*Δ and *vps1*Δ *dnm1*Δ, mutants defective in peroxisome fission. Since Vps1, but not Dnm1, is required for fission in glucose-grown cells, single and double mutants of Vps1 show a reduced basal number of peroxisomes. d) GLM analyses showing that *vps1*Δ and *vps1*Δ *dnm1*Δ show significantly reduced peroxisome proliferation in response to tunicamycin compared to WT. The analyses also shows that tunicamycin treatment significantly reduces the cells without peroxisomes in *vps1*Δ and *vps1*Δ *dnm1*Δ suggesting faster *de novo* biogenesis in cells which lacked peroxisomes. e) Cartoon showing the peroxisome inheritance defects in *inp1*Δ and *inp2*Δ mutants. f) Quantification of proportion of cells without peroxisomes in the inheritance mutants 5h after treatment with DMSO or 0.5µg/mL tunicamycin. Bars indicate an average of two experiments, and error bars indicate standard deviation. Statistical analyses performed by comparing tunicamycin treated samples vs. DMSO.

We next tested whether the increase in peroxisome number during ER stress depended on peroxisome division or on their *de novo* synthesis. Vps1 is the primary mediator of peroxisome fission in cells grown in presence of glucose ^68^, and its deletion resulted in only a partial reduction in tunicamycin-driven peroxisome proliferation compared to WT cells (Figure 6c,d). We also quantified this tunicamycin response in *dnm1*Δ single and *vps1*Δ *dnm1*Δ double mutants. While the loss of *DNM1* alone had an insignificant effect, the absence of both *DNM1* and *VPS1* significantly, but still only partially, impaired peroxisome proliferation after tunicamycin treatment (Figure 6c,d), suggesting additional peroxisome formation occurs by the *de novo* pathway. In DMSO-treated cells, the median number of peroxisomes was 1 in both *vps1*Δ and *vps1*Δ *dnm1*Δ mutants, while tunicamycin-treated cells had a median number of 2 peroxisomes per cell. This suggests that tunicamycin treatment led to the *de novo* formation of at least one additional peroxisome. In comparison, WT cells had a median number of 3 peroxisomes in DMSO treatment, and 4.5 peroxisomes after tunicamycin treatment. Thus, of the ∼1.5 additional peroxisomes formed in WT in response to tunicamycin, at least one could be attributed to *de novo* biogenesis. Overall, our results demonstrate that *de novo* biogenesis accounts for approximately two-thirds of the peroxisome formation in response to ER stress.

Further evidence supporting the role of *de novo* biogenesis in increased peroxisome production during ER stress comes from analyzing the proportion of cells containing peroxisomes in the inheritance mutants *inp2*Δ and *inp1*Δ (Figure 6e-f). In the absence of Inp2, most daughter cells fail to inherit peroxisomes from mother cells and therefore rely exclusively on *de novo* biogenesis to produce their first peroxisome ^69,70^. Conversely, most mother cells fail to retain peroxisomes in the absence of Inp1 ^71^, resulting in the total transfer of peroxisomes to their daughter cells. We observed ∼45% of *inp2*Δ cells lacked peroxisomes under DMSO-treated conditions; however, tunicamycin treatment reduced this proportion to ∼22% (Figure 6f), strongly suggesting that ER stress stimulated *de novo* biogenesis. Additionally, we found ∼25% of *inp1*Δ cells lacked peroxisomes after tunicamycin treatment, compared to ∼40% under DMSO treatment conditions (Figure 6f). Notably, the timeline of peroxisome proliferation after tunicamycin treatment observed in our study is typical of *de novo* peroxisome formation (∼4-5 h) ^68,69,72^, further corroborating the dominant contribution of the *de novo* pathway to this response.

## DISCUSSION

### ER stress induces peroxisome proliferation in an evolutionarily-conserved manner

Most studies of peroxisome induction have focused on the transcriptional regulation of peroxisome biogenesis in response to nutritional cues; one well-studied condition involves the stimulation of peroxisome biogenesis with oleate ^10,73-76^, which necessitates the induction of peroxisomes and their contents to metabolize this fatty acid. Few studies have focused on the effects of environmental cues, such as various forms of stress (e.g., misfolded or unfolded proteins, salt, heat, lipid, or oxidative stress), on peroxisomes ^19-22^. Moreover, these studies lack a comprehensive analysis of the underlying mechanisms and signaling pathways involving peroxisome biogenesis and proliferation (Figure 7).

**Figure 7:**
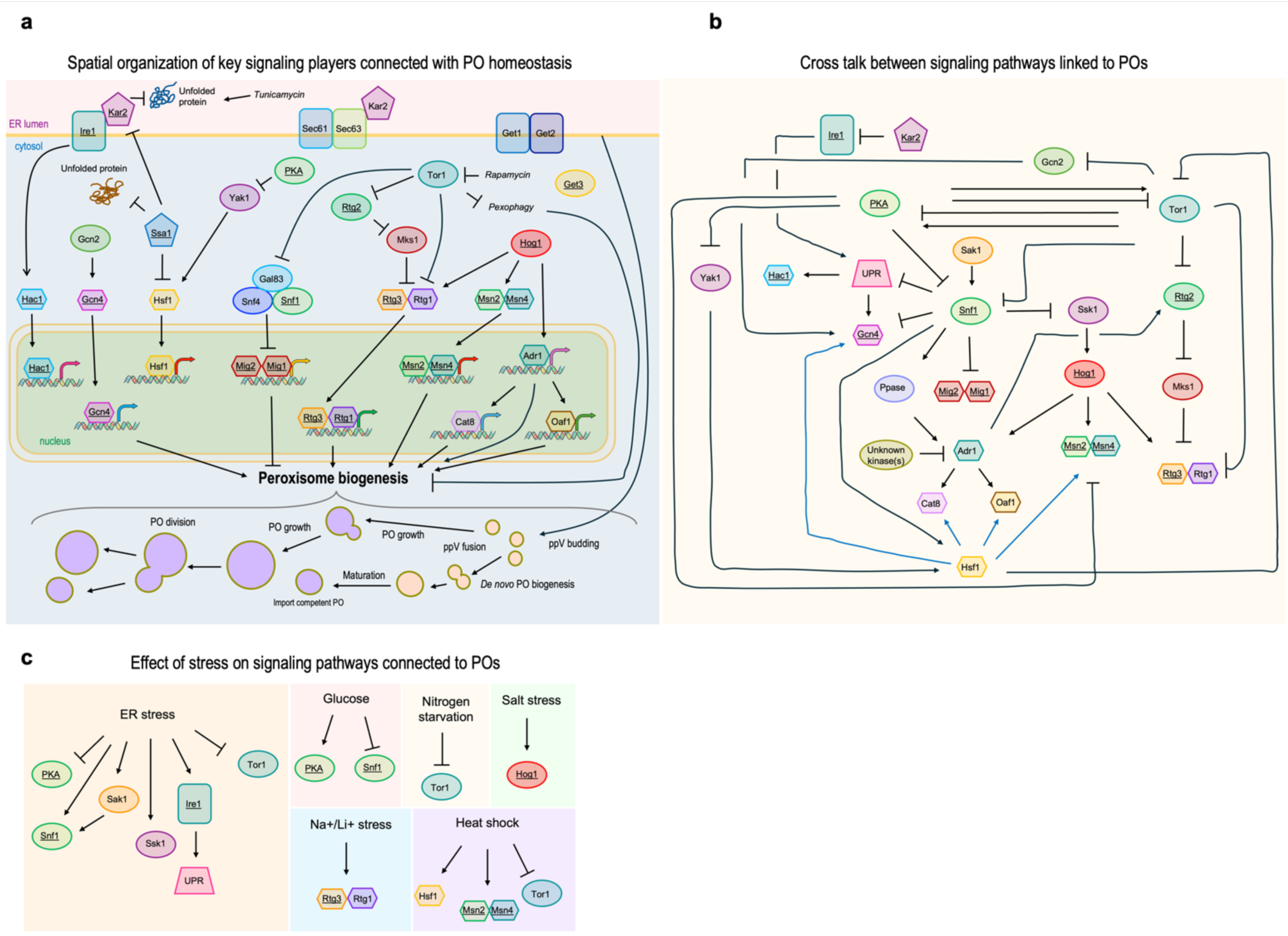
Summary of *S. cerevisiae* signaling pathways implicated in the induction of peroxisomal genes. a) Cartoon explaining the spatial organization of key signaling players that are activated or inhibited during ER stress and have been linked with peroxisome formation, maintenance or function previously. Players studied here using mutants or drug treatments are underlined. b) Overview of the cross talk between signaling pathways shown in (a). c) Overview of the impact of different stresses on key signaling players tested in this study based on previous reports. References for these pathways are as follows: a) Ire1 activation of Hac1 ^125^, Gcn2 activation of Gcn4 ^126^, PKA inhibition of Yak1 ^127^, Yak1 activation of Hsf1 ^127^, Ssa1 prevention of cytosolic unfolded protein accumulation and inhibition of Hsf1 ^30^ and Kar2 ^128^, Hsf1 activation of gene expression ^129^, Gcn4 activation of gene expression ^45,46^, Tor1 inhibition of Snf1 ^130^ and pexophagy ^131^, Snf1 inhibition of Mig1/2 ^64,132^, Mig1/2 repression of peroxisome biogenesis ^133^, Tor1 regulation of Mks1 via Rtg2 ^134^, Mks1 inhibition of Rtg1/3 ^134^, Rtg1/3 induction of peroxisome biogenesis ^86^, Tor1 inhibition of Rtg1/3 ^134^, Tor1 inhibition by rapamycin ^135^, Hog1 activation of Rtg1/3 ^136^ and Msn2/4 ^137^, Msn2/4 requirement for peroxisome biogenesis ^17^, Hog1 activation of Adr1 ^22^, Adr1 induction of peroxisome biogenesis ^73^, via Cat8 and Oaf1 ^138^. b) Kar2 inhibition of Ire1 ^42^, Ire1 activation of UPR ^125^, Tor1 inhibition of Gcn2 ^139^, Rtg1/3 ^140^, Rtg2 ^134^ and Snf1 ^130^, PKA and Tor1 cross regulation ^141^, PKA inhibition of Yak1 ^127^, Snf1 ^96^ and Msn2/4 ^142^, Yak1 activation of Hsf1 ^127^, Hsf1 inhibition of Tor1 ^29^, Hsf1oe activation of Gcn4 ^143^, as well as Cat8, Oaf1 and Msn2/4 ^54^, UPR activation of Hac1 ^125^ and Gcn4 ^46^, Sak1 activation of Snf1 ^144^, Snf1 inhibition of UPR ^97^, Gcn4 ^143^, Mig1/2 ^64,145^ and Ssk1 ^93^, Snf1 activation of Adr1, via a phosphatase (PPase) ^94,146^, as well as Hsf1 ^129^, Unknown kinase activating Adr1 ^146^, Ssk1 activation of Hog1 ^147^, Adr1 activation of peroxisome biogenesis via Cat8 and Oaf1 ^75^, and of Rtg2 ^87^, Rtg2 inhibition of Mks1 ^87,148^, Mks1 inhibition of Rtg1/3 ^87,148^, Hog1 activation of Adr1 ^22^, Msn2/4 ^137^ and Rtg1/3 ^136^. c) ER stress inhibition of PKA ^61^ and Tor1 ^149^, ER stress activation of Snf1 ^93^, Sak1 ^144^, Ssk1 ^150^ and Ire1 ^151,152^, Sak1 activation of Snf1 ^144^. Glucose activation of PKA ^141^ and inhibition of Snf1 ^153^. Nitrogen inhibition of Tor1 ^154^, salt stress activation of Hog1 ^22^, Na^+^/Li^+^ stress activation of RTG pathway ^22^, Heat shock activation of Hsf1 ^155^ and Msn2/4 ^54^, and inhibition of Tor1 ^29^.

Our study reveals that ER stress induces peroxisome proliferation in S. cerevisiae (Figure 1), in *K. phaffii* (Figure 5c,d), which diverged from *S. cerevisiae* ∼250 M years ago ^77^, as well in human cells, representing ∼1 billion years of divergence from *S. cerevisiae* (Figure 5g). We show that peroxisomes are required to respond to ER stress (Figure 5f), a normal condition yeast cells experience outside of typical laboratory conditions, regardless of carbon source. Thus, our results demonstrate that peroxisomes play a central role in the conserved, cellular adaptation to ER stress. Additionally, our findings also dispel the misconception that peroxisomes are dispensable when yeast cells metabolize glucose.

Previously, we showed that peroxisome abundance, proliferation and function are integrated into many diverse processes and are influenced by interactions and signaling from multiple subcellular compartments including the mitochondria, nucleus and cytosol ^36^. Additionally, vacuole-mediated pexophagy regulates peroxisome turnover ^3,36^. While peroxisomes can arise by budding of vesicles from the ER, with ER supplying lipids for peroxisome growth via contact sites ^37,78,79^, this study extends the influence of ER and the cytosol on peroxisome homeostasis, further emphasizing the interconnectivity and coordination between compartments in mounting an appropriate response to stress and maintaining cellular homeostasis.

### Tunicamycin-induced peroxisome proliferation is primarily by *de novo* biogenesis alongside growth and division

Previous studies on *de novo* peroxisome biogenesis have shown that peroxisomes arise in about 4-5 h ^72^. Our analysis of peroxisome number performed within the same timeline after ER stress-induction primarily reflects increases due to *de novo* peroxisome biogenesis with a smaller contribution (∼one-third) from growth and division. Reduced pexophagy cannot account for the increase in peroxisomes because *atg36*11 cells, which are blocked in this process ^67^, also show significant tunicamycin-induced peroxisomes (Figure 6b).

The significance of *de novo* peroxisome biogenesis remains a mystery because growth and division is the dominant pathway for generating new peroxisomes in glucose-grown yeast cells ^69^. Since our experimental strategy uses a basal state where yeast cells cannot synthesize new peroxisomes by growth and division (*vps1*Δ or *vps1*Δ *dnm1*Δ cells), any new peroxisome generated in response to tunicamycin treatment must arise via *de novo* biogenesis from the ER. In glucose-grown cells, previous studies have shown that peroxisome division mainly involves Vps1, whereas Dnm1 is required for peroxisome division in oleate ^68^. Accordingly, while *vps1*Δ and *vps1*Δ *dnm*1Δ were partially defective in tunicamycin-induced peroxisome proliferation, whereas *dnm1*Δ cells did not show any significant difference compared to WT in their tunicamycin-response. Overall, these results show that both mechanisms of peroxisome production contribute to tunicamycin-driven increase in peroxisomes (Figure 6c-d). Additional evidence for tunicamycin-induced *de novo* peroxisome formation comes from *inp1*11 and *inp2*11 cells, which have peroxisomes only in daughter and mother cells, respectively, due to inheritance defects ^70^, and new peroxisomes in mother and daughter cells in these respective mutants can only have arisen *de novo* ^80^ (Figure 6e-f).

### Sources of stress and pathways causing peroxisome proliferation

ER stress is triggered by many environmental conditions and arises from multiple factors including misfolded and unfolded proteins, redox stress, lipid bilayer stress, or mistargeting of ER proteins ^81^. We show that activation of the UPR, triggered by activation of Ire1 kinase, Hac1 and Gcn4 is dispensable for peroxisome proliferation (Figure 2a-c). Similarly, lipid bilayer stress caused by inositol deprivation (Figure 2d-e) ^47,82^ and activation of the ER Stress Surveillance pathway (Figure 2f-g) ^48^ also do not trigger peroxisome induction. Instead, our results point to the mistargeting of ER proteins, as seen in *get3*11 cells, and/or the subsequent accumulation of misfolded proteins in the cytosol, as observed in *ssa1*11 cells, as primary triggers of peroxisome proliferation (Figure 3b,c). Our findings thus highlight a novel mechanism linking protein quality control and cytosolic stress to peroxisome biogenesis.

The activation of peroxisome proliferation by rapamycin (via Tor1 inhibition), either partially in *S. cerevisiae* and human cells or substantially in *K. phaffii*, shows for the first time the role of active Tor1 in negatively regulating peroxisome proliferation (Figure 5a-d, 5g). Consistent with this, Tor1 is inactivated during ER stress ^60^ (Supplemental Figure 4). Surprisingly, activation of the cytosolic heat shock response in *ssa1*11 cells also increased the basal number of peroxisomes, and while this effect was further amplified by tunicamycin treatment, the increase was modest in comparison to WT (Figure 3c,g). This suggests that the cytosolic heat shock response is also a potent regulator of peroxisome biogenesis, and that it partially accounts for tunicamycin-driven peroxisome proliferation. Notably, ER stress can also indirectly activate Hsf1 through the inhibition of PKA and in principle, the activation of Snf1 (Figure 7) ^83^. We propose Hsf1 activation plays a dual role: it both alleviates ER stress ^43,84^ and directly promotes peroxisome biogenesis. Hsf1 can activate Oaf1, Cat8 and Msn2/4 ^54^, which control expression of genes involved in peroxisome proliferation ^85^. Additionally, Hsf1 can also inhibit Tor1 ^29^, which in turn can activate the RTG pathway ^62,86,87^, PKA ^88^ and Gcn4 ^45^ (Figure 7a-b).

It is reasonable to wonder whether it is reductive stress that is common between the induction of ER stress and cytosolic misfolded protein pathways, both of which cause peroxisome induction. One compelling possibility is that altered reduced/oxidized ratios of thioredoxin (TRX) or glutathione (GSH), in response to these stresses, may underlie peroxisome induction. Supporting this idea, the constitutive activation of the UPR in yeast increases the GSH/GSSG ratio, leading to a more reduced state in both the ER and cytosol of yeast ^89^. Similarly, a more reduced redox state of the cytosol, but not of the ER, is observed during oxidative protein folding in the ER without UPR induction ^89^. Indeed, we show that yeast cells subjected to reductive stress initiated by DTT treatment significantly induced peroxisomes (Figure 1g).

However, because the TRX pathway predominates in the yeast cytosol over the backup GSH pathway, and only perturbations of the former activate Hsf1 ^90^, redox balance of one or both pathways could play a role in peroxisome homeostasis. Yeast peroxisomes have several enzymes involved in these two pathways such as Ahp1, Gpx1, catalase and glutathione-S-transferases ^91^. These findings suggest that redox imbalance, whether from altered TRX or GSH pathways, may signal integrating stress responses to regulate peroxisome homeostasis and adaptation.

### Contribution of other signaling pathways in tunicamycin-induced peroxisome proliferation

While the Snf1 pathway and its downstream effectors (e.g., Adr1, Oaf1 and Cat8) ^44,92^ regulate peroxisome biogenesis in yeast metabolizing oleate ^10,18^, and ER stress has been shown to induce Snf1 ^93^, our data show that *snf1*11 cells grown in SD medium have similar peroxisome numbers as WT cells (Figure 4c). Moreover, under our assay conditions, we did not observe Snf1 activation, corroborating our conclusion that tunicamycin-driven peroxisome proliferation is Snf1-independent (Supplemental Figure 4, Figure 4c). Additionally, activated Snf1 would be expected to relieve the repression of peroxisome biogenesis by phosphorylation and nuclear export of the transcriptional repressors Mig1 and Mig2 ^64^, but both *mig1*11 and *mig2*11 cells had peroxisome numbers comparable to WT cells under basal and tunicamycin-induced conditions (Figure 4e). This result is consistent with the high degree of pathway redundancy involved in peroxisome biogenesis ^10,16,18,45,73,94,95^ and the extensive crosstalk between Snf1 and other signaling pathways (Figure 7b) ^92,93,96-99^.

Similarly, although the RTG pathway has been reported to induce peroxisome biogenesis during salt stress ^22^ and partially upon tunicamycin treatment ^20^, and Hog1 is required for fatty acid metabolism during salt stress and oleate induction ^22^, our results suggest a different mechanism under ER stress. Our data show *hog1*11, *rtg1*11, *rtg2*11, *rtg3*11 and *msn2*11*msn4*11 mutant cells exhibited basal peroxisome levels comparable to those of WT cells, with tunicamycin-induced peroxisome proliferation also mirroring the WT response (Figure 4b-d). These findings suggest that the mechanism by which peroxisomes are induced during salt stress differs from that we observe during tunicamycin-induced proliferation, highlighting the varied mechanisms regulating peroxisome biogenesis in response to distinct stressors. While our data indicates these pathways to individually be dispensable for tunicamycin-mediated peroxisome proliferation, they could potentially be redundant. Additionally, ER stress inhibits PKA ^61^. Our results indicate that the activation of the PKA pathway, using TPK1oe ^65,66^, impaired tunicamycin-induced peroxisome proliferation compared to WT cells (Figure 4f-g), and we observed altered PKA activity after 4h (Supplemental Figure 4). This supports the idea that tunicamycin-induced peroxisome proliferation partially involves mechanisms dependent on PKA activity. While both PKA and Tor1 are inhibited during ER stress, the complex interplay between these two kinases in the context of ER stress has not been worked out yet, so it will be particularly exciting to understand how their mutual relationship impinges on peroxisome homeostasis. Overall, the insignificant or partial effects of the signaling pathway mutants on peroxisome biogenesis can be attributed to the possible overlap in their target genes and would also explain why these mutant genes have not been uncovered in other genetic screens as candidates implicated in peroxisome biogenesis.

### The physiological relevance of peroxisomes in the cellular adaptation to ER stress

Our data indicate that peroxisome biogenesis is critical for cell survival following induction of ER stress using tunicamycin. This suggests that peroxisomes support specific metabolic adaptations critical for cellular survival during stress. Cellular ER stress in yeast shifts metabolism away from glycolysis/fermentation ^100^ to lipolysis (for fatty acids), respiration (for ATP), glyoxylate pathway (for citrate entry into TCA), gluconeogenesis (for glucose-6-P in the ER), the pentose phosphate pathway (for NADPH and redox balance) and acetyl-CoA production by peroxisomal fatty acid oxidation for mitochondrial ATP production ^20,22^. The production of acetyl-CoA and NADH to feed the TCA cycle is typically through fatty acid oxidation in peroxisomes. Consistent with this, we find that Pot1, a key enzyme in fatty-acid oxidation, but which is repressed ordinarily by glucose, is induced by tunicamycin (Figure 1h-i). Other studies have shown the induction of other enzymes involved in fatty-acid oxidation and transport by DTT and tunicamycin ^101,102^. Thus, ER stress not only induces peroxisomes, but reprograms them to enable cellular adaptation.

Since the peroxisome proliferation we see is in glucose-grown cells, the cellular source of fatty acids required for acetyl CoA and NADH production remains unclear. We speculate that lipid droplets (LDs), which can produce free fatty acids either via lipolysis and/or lipophagy could be a potential source ^103-106^. Notably, ER stress in yeast induces LD formation ^107^, as well as lipolysis, especially via microlipophagy ^104^, which we hypothesize as the source of fatty acids for peroxisomal β-oxidation.

ER stress and tunicamycin cause ER membrane expansion ^108^, which would also be required for the generation of pre-peroxisomal vesicles by *de novo* biogenesis from the ER ^109^. This membrane expansion requires lipid (especially phospholipids) synthesis and helps alleviate ER stress ^108^. The hydrolysis of triacylglycerols (TAG) and steryl esters (STE) stored in LDs by TAG lipases and STE hydrolases ^110,111^ could also provide fatty acids and sterols for such membrane expansion.

Several lines of evidence suggest that peroxisomes play an important role in cellular adaptation to ER stress. First, the conservation of peroxisome induction in response to ER stress points strongly to an evolutionarily-important pathway. Second, yeast *pex3*11 cells, which lack functional peroxisomes, and *pex34*11 cells that accumulate ROS ^20^, exhibit significantly reduced ability to adapt to ERS when pre-exposed to a low dose, followed by a high dose of tunicamycin (Figure 5f). Third, peroxisomes provide metabolic support for mitochondrial energy production by providing acetyl-CoA, which requires a functional peroxisomal fatty-acid β-oxidation pathway, and NADH ^112^. Cells lacking peroxisomal β-oxidation cannot significantly expand their maximum oxygen consumption rate after tunicamycin treatment, unlike WT cells ^20^. Fourth, peroxisomes help reduce ROS accumulation during ER stress responses ^20^. Finally, peroxisomes are involved in lipid metabolism, whose disruption impacts lipid homeostasis, which in turn, aggravates ER stress ^113,114^.

### Potential applications of the induction of *de novo* peroxisome biogenesis pathways in disease states

Peroxisomes are essential for cellular function, and their disruption can lead to severe or lethal consequences. However, milder cases, often linked to temperature-sensitive missense mutations in key *PEX* genes, can be managed with appropriate treatments ^27^. Among the pharmacological treatments being considered are those designed to increase peroxisomes by either inducing peroxisome biogenesis and/or reducing pexophagy ^115^. While 4-PBA induces peroxisome proliferation and improves biochemical function (very long chain fatty acid beta-oxidation rates and very long chain fatty acid and plasmalogens levels) in fibroblast cell lines from patients with milder PBD phenotypes ^116^, it is mechanistically diverse and peroxisomal proliferators that have been discovered to more specifically induce peroxisomes in rodents, do not induce human peroxisomes ^13^. Our studies provide several additional ways in which peroxisome numbers might be induced, including ER and reductive stress, tunicamycin, activation of heat shock response and Tor1 inhibition. Importantly, the induction of *de novo* biogenesis pathways might alleviate the symptoms of patients impaired in the peroxisome division machinery (e.g. PEX11 ^117^). Finally, because there are many human diseases associated with an activation of the ER stress response ^118,119^, and the cellular adaptability to ER stress declines with age, infections and environmental exposures ^120,121^, the role of peroxisomes in alleviating these processes will be particularly exciting to pursue further.

## AUTHOR CONTRIBUTIONS

NS and SS co-conceived the study, interpreted data, and co-wrote the first draft. NS generated strains and plasmids, designed and performed experiments and analyzed data. MN performed GLM analyses. JF discovered the role of Tor1 inhibition on peroxisome number, designed and performed experiments in *K. phaffii*. FM and LM performed human cell experiments. LT and NS performed experiments on UPRE-GFP and HSE-GFP. TS assisted in strain generation. JA and SS supervised the study and obtained funding. NS, MN, JC, FM, JA and SS edited the manuscript.

## ACKNOWLEDGEMENTS

This study was funded by grant NIDDK 41737 to SS and JA. SS is a Tata Chancellor Professor in Molecular Biology. NS is grateful for partial support from Molecular Biology Cancer Fellowship (UC San Diego). FM is the recipient of a career development award from Seattle Children’s Research Institute. We are thankful to Qihao Liu and Lorraine Pillus (UC San Diego) for constructing the *msn2*Δ*msn4*Δ strain, and Haley Schultz for the sHS60 strain. We are also grateful to the following colleagues for providing strains and plasmids: Brenda Andrews and Charlie Boone (Univ. of Toronto) for the *kar2-159*-strain; Paul Herman (The Ohio State University) for pPHY2056 and pRS426 plasmids; Randolph Hampton (UC San Diego) for providing pRH1209 plasmid; Richard Rachubinski (U. of Alberta) for the Pot1-GFP strain; Trey Ideker (UC San Diego) for BY4741 strain. We gratefully acknowledge the support of Andrew Longenecker and the PBD project for providing the human primary fibroblasts used in this study.

## MATERIALS AND METHODS

### Strains and Plasmids

Yeast strains were generated using lithium acetate-PEG-mediated transformation and are listed in Supplemental Table 1. Strains expressing P*_TDH3_*-GFP-ePTS1-T*_PGK1_*, (i.e., GFP-ePTS1) 4xUPRE-P*_CYC1_*-GFP-T*_ACT1_* (i.e., UPRE-GFP) or 4xHSE-P*_CYC1_*-GFP-T*_ACT1_* (i.e., HSE-GFP) were generated by transforming linear constructs obtained by double digesting the plasmids pNS31, pNS32 or pLT1, respectively, with PciI and NotI. All three constructs were targeted after the STOP codon of the *UBC9* ORF. Except where noted, parental knock-out strains were obtained from the MATα deletion library (BY4742) ^122^. *INP2* and *DNM1* genes were knocked out by transforming the corresponding deletion constructs, which in turn, were generated by fusing hphMX6 or Zeo cassettes on either side with ∼600-1000bp sequences upstream and downstream of the respective ORFs. 4xUPRE-P*_CYC1_*-GFP-T*_ACT1_* was amplified from pRH1209 ^123^. For generating strains overexpressing TPK1 (TPK1oe), plasmid pPHY2056 ^65^ was transformed in yeast; the corresponding control strain was made by transforming the empty vector (EV), pRS426 in yeast.

### Yeast growth, treatments, microscopy and Western blotting

Except when noted, *S. cerevisiae*, log phase cells grown in synthetic defined (SD; 6.7g/L Yeast nitrogen base (YNB) + 0.79g CSM) + 2% dextrose at 30°C and shaken at 250 RPM were used. For experiments with HSE-GFP and *kar2-159*, cultures were grown at 25°C and 250 RPM. Cultures were resuspended in fresh media at an OD∼0.05-0.1 prior to performing temperature shifts (to 30°C or 37°C for 3h as indicated in figure legends) or treatments with tunicamycin (Sigma, T7765; 1mg/ml stock in DMSO) or rapamycin (Sigma, R0395; 1mg/ml stock in DMSO) or phytosphingosine (Sigma, Cas No. 554-62-1; 10mM stock in DMSO) or DMSO or DTT (Roche, 10708984001; 1M stock in H_2_O). TPK1 overexpression experiments were performed in SD-Ura media to maintain selection for the plasmids. For all the experiments with *K. phaffii,* cell growth and treatments were done in YPD media (10g/mL Yeast extract + 20g/mL Bactopeptone + 2% Dextrose). SD-Ino media was prepared using YNB without inositol (US Biological Life Sciences; Y2030-01).

Cells were imaged using a Zeiss Plan Apochromat 60x/1.4 Oil DIC objective mounted on an Axioskop 2 mot plus microscope (Zeiss) equipped with Axio Cam HRm camera and HBO 100 mercury lamp. For 3D imaging to count the number of peroxisomes per cell, Z stacks consisting of 8 slices (for *S. cerevisiae*) or 6 slices (for *K. phaffii)* were acquired with a Z spacing of 1µm. For UPRE-GFP, HSE-GFP and Pot1-GFP, only a single slice at the focal plane was imaged. Exposure times were kept identical across all experiments for each fluorescent marker, whereas DIC images were acquired with auto exposure.

Western blotting was performed from cell extracts generated after TCA-precipitation using the following antibodies: Anti-ScActin Rabbit pAb, Gift from Michael Yaffe (1986); Phospho-S6 Ribosomal Protein (Ser235/236) (D57.2.2E) XP Rabbit mAb, Cell Signaling Technology; #4858; Phospho-AMPKα (Thr172) (40H9) Rabbit mAb, Cell Signaling Technology; #2535; Phospho-p38 MAPK (Thr180/Tyr182) (D3F9) XP Rabbit mAb, Cell Signaling Technology; #4511; Phospho-PKA Substrate (RRXS*/T*) (100G7E) Rabbit mAb, Cell Signaling Technology; #9624; Goat Anti-Rabbit IgG (H+L)-HRP Conjugate, Bio-RAD; #1721019.

### Human fibroblast cell culture and microscopy

Human primary fibroblasts were cultured at 37°C with 5% CO₂ in high-glucose Dulbecco’s Modified Eagle Medium (DMEM; Gibco) supplemented with 15% (v/v) heat-inactivated fetal bovine serum (FBS; VWR). All experiments were conducted using fibroblasts with fewer than 10 passages. For imaging, fibroblasts were seeded in glass-bottom 96-well plates (Cellvis) and treated with the indicated concentrations of tunicamycin (Sigma), rapamycin (Sigma), or 4-phenyl butyric acid (4-PBA) (Sigma). After a 12-hour incubation, cells were fixed with formaldehyde and stained with DAPI (Thermo Fisher Scientific), PMP70-AF488 (anti-PMP70 antibody [EPR5614]: abcam; Thermo Fisher Scientific), and Phalloidin-AF750 (Thermo Fisher Scientific). Each treatment condition included three biological replicates. For imaging, 6–8 3D confocal z-stacks per well were acquired using a Zeiss 980 confocal microscope equipped with a 63×/1.2 NA water immersion objective lens and NIR detector. Each z-stack had dimensions of 4096 × 4096 × 31 voxels, corresponding to a physical volume of approximately 210 × 210 × 6 µm³.

### Data analyses and Graphing

Figure images for depicting the number of peroxisomes in cells were made by generating maximum intensity Z projections for peroxisome channel (GFP) and average intensity Z projection for cell outline (DIC) channel. Figure images for depicting UPRE-GFP, Pot1-GFP and HSE-GFP were made with single-Z slice images. Identical brightness and contrast settings were applied across all images within a figure panel to ensure fair comparisons, using FIJI/ImageJ software. Scale bars in all figures represent 5 µm, except for the human fibroblasts where a 50 µm scale bar was used. Quantification for number of peroxisomes per cell (for GFP-ePTS1 and Pex3-GFP) as well as for intensity of GFP signal per cell (for UPRE-GFP and HSE-GFP) was performed using the perox-per-cell software ^38^ using a minimum peroxisome area threshold of 1 pixel and the following values for the software’s peroxisome detection sensitivity: 0.0064 for GFP-ePTS1, 4xUPRE-GFP, and 4xHSE-GFP; 0.003 for Pex3-GFP. For quantification of Pot1-GFP, UPRE-GFP and HSE-GFP levels, the values for GFP signal intensities for each cell were divided by 10,000x the value of their respective cell areas to obtain the ‘normalized GFP signal’ used in the plots. GraphPad prism and Microsoft excel were used to generate the graphs. All yeast experiments were done twice, in separate experimental batches on different days. A batch consisted of a group of 1-4 mutants that were tested together with a common WT control. Two different batches for a mutant thus represented two biological replicates, each with at least a hundred cells. While the graphs in the figures depict data from a single replicate, GLM analyses were performed on data pooled across batches and thus included both replicates, to rule out discrepancies due to batch-specific effects. The whiskers in the box plots represent 1-25 and 75-99 percentiles of the data, whereas the top and bottom 1 percentile are represented as dots outside these bounds. In the histograms, the X-axis was cropped mostly at 15 or sometimes at 20 peroxisomes/cell, since any higher bin values accounted for <1% of the frequencies. We observed significant cell death for *kar2-159* ts mutant cells at 37°C, and for *hac1*Δ after tunicamycin treatment, therefore we manually excluded dead cells from the histograms for those experiments. For this, any atypical cell that showed shriveling, presence of large vacuoles, or obvious damage and abnormal shape as visualized on the DIC channel was marked as dead. Our empirical observations indicated that almost all cells with zero peroxisomes were dead or severely damaged, except in *inp1*Δ, *inp2*Δ, and *vps1*Δ *inp2*Δ, where the lack of peroxisomes can be attributed to inheritance defects. We also did not exclude dead cells from the data for GLM analyses (see below), hence many of the mutants that appear to have significant changes in zero peroxisome counts are those where we observed more death after tunicamycin treatment (viz. *hac*1Δ, *hog1*Δ). Using GraphPad prism, after confirming that the data did not fit normal distribution using D’Agostino & Pearson, Anderson-Darling, Shapiro-Wilk, and Kolmogorov-Smirnov tests, we used the Mann-Whitney test to evaluate statistical significance between control and treatment groups. Welch’s t test was used to evaluate statistical significance for assays testing viability of *pex3*Δ vs WT after tunicamycin treatment and to compare the number of cells lacking peroxisomes in *inp1*Δ and *inp2*Δ vs WT (n=2 experiments).

### Modeling tunicamycin-induced peroxisome counts across mutant strains

We used generalized linear modeling (GLM) in R (version 4.4.0) to model the distribution of peroxisome (PO) counts in cells across treatment conditions and strains. Due to the presence of a high number of cells with zero counts in some strains, we modeled distributions with hurdle models ^55^ using the “pscl” R package (version 1.5.9), where zero and non-zero (positive) peroxisome counts are captured using separate functions. A binomial distribution was used to model zero counts, and a negative binomial distribution was used to model non-zero counts. We found that, based on the Akaike Information Criterion, hurdle models fit the data better than zero-inflation and simple negative binomial models.

To compare a mutant strain’s tunicamycin response to WT, we first combined that strain’s peroxisome count data for both the DMSO and tunicamycin treatment conditions with the full set of WT count data collected across all experiments and strains. This included data from both DMSO- and tunicamycin-treated WT cells. We then used GLM to fit the data using a model formula that included strain, treatment, experimental batch, and an interaction term to quantify strain-specific responses to tunicamycin.

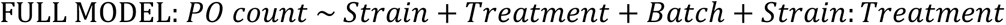

We tested whether the mutant strain’s response to tunicamycin was significantly different than WT by performing a likelihood ratio test where GLM results using the following reduced model were compared to those from the full model.

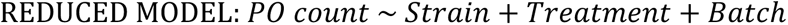

*P*-values collected from these tests across strains were then adjusted for the false discovery rate using the Benjamini-Hochberg method ^124^. We considered mutant strains with *P*-values < 0.05 as those with tunicamycin responses that were significantly different from WT.

To determine whether a mutant strain’s response to tunicamycin was stronger or weaker compared to WT, we examined the value of the coefficients for the *Strain:Treatment* interaction term in the mutant strain’s full GLM model. Because hurdle models have two modeling components, there are two interaction term coefficients to consider: one from the zero-count model and one from the non-zero count model. For heatmap visualization, the coefficients from a given hurdle modeling component were z-score normalized across the values obtained from all strains (i.e., normalized by column for the heatmaps). For consistency, positive values in these visualizations indicate that the modeling component predicts a stronger tunicamycin response in a mutant strain compared to WT. That is, it predicts a response resulting in higher peroxisome counts. As mentioned earlier, we observed a significantly large proportion of dead cells in *hac1*Δ and *hog1*Δ strains following tunicamycin treatment. The zero-count model interaction term coefficients in the GLM analyses on those strains reflected this, resulting in higher predicted zero-count instances. To ensure that the presence of dead cells did not bias our analyses, we made inferences on the relative effects of tunicamycin-induced peroxisome proliferation for mutants vs. WT based only on the interaction term coefficients of the positive-count model.

**Supplemental Figure 1:**
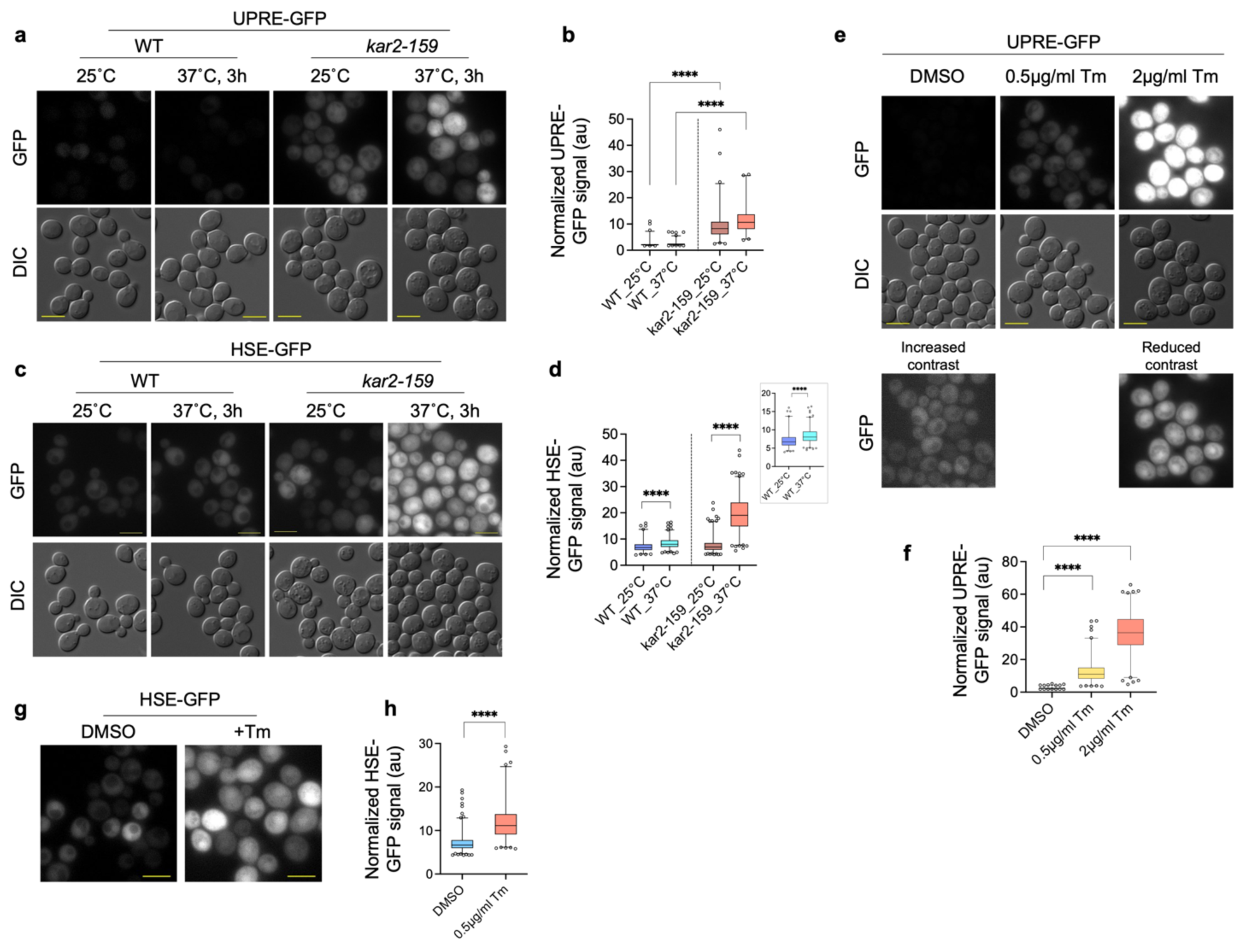
Protein misfolding defect induces UPR and heat shock response. a-d) Single Z-slice images (a,c) and quantification (b,d) showing UPRE-GFP (a-b) or HSE-GFP (c-d) levels per cell in WT and *kar2-159* cells after growth at 25°C and after 3h of growth at 37°C. Inset in (d) to show WT at different temperatures [(b) N_cells_ for WT: 25°C: 337, 37°C: 594; N_cells_ for *kar2-159*: 25°C: 362, 37°C: 231; (d) N_cells_ for WT (same data as Figure 3e): 25°C: 416, 37°C: 620; N_cells_ for *kar2-159*: 25°C: 863, 37°C: 795]. e-h) Single-Z-slice images (e,g) and quantification (f,h) of UPRE-GFP and HSE-GFP levels in WT cells after treatment with either tunicamycin (Tm) or DMSO [(f) N_cells_: DMSO: 786, 0.5µg/ml Tm: 531, 2µg/ml Tm: 532; (h) N_cells_: DMSO: 819, 0.5µg/ml Tm: 546]. Scale bar: 5µm.

**Supplemental Figure 2:**
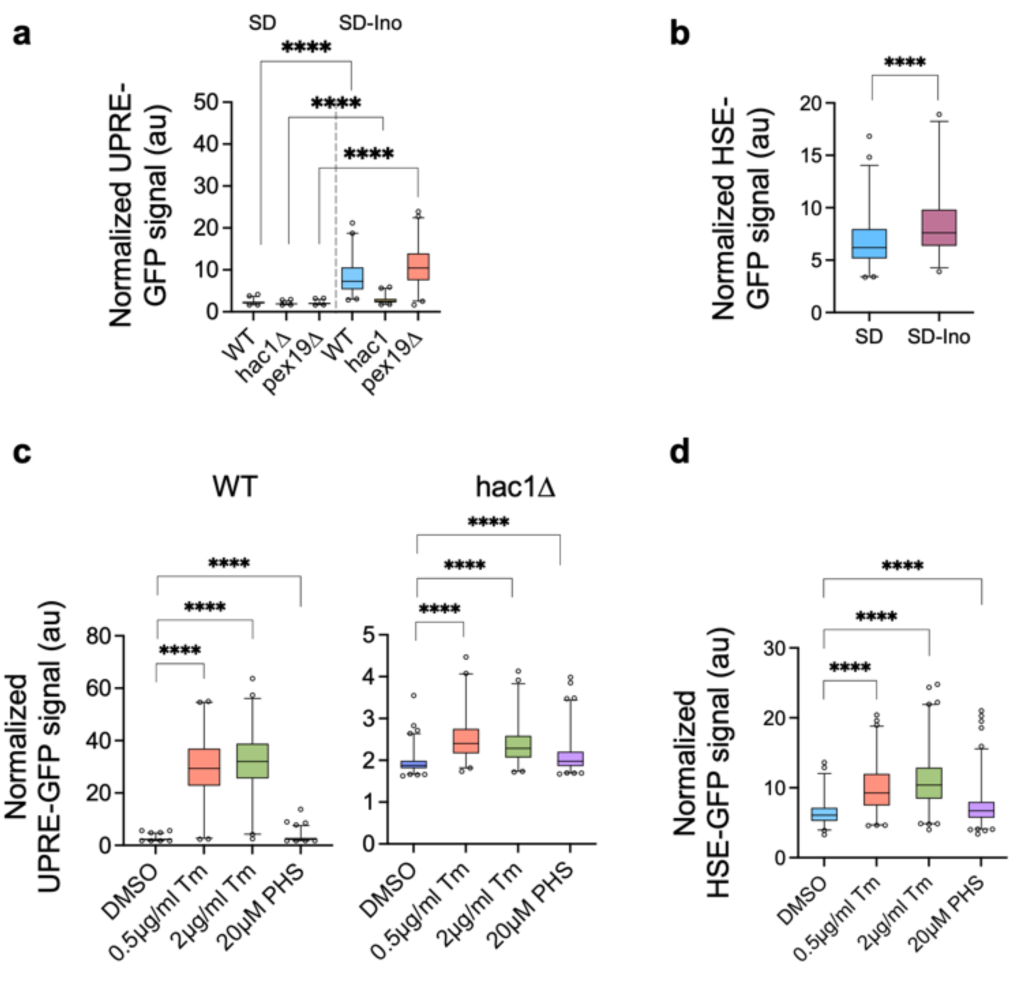
Lipid stress or activation of ERSU do not induce UPR or heat shock response. a) Inhibition of peroxisome formation does not impair UPR activation as indicated by the increase in UPRE-GFP signal in *pex19*Δ. *hac1*Δ cells served as a negative control; the basal increase in the GFP signal in *hac1*Δ, although significant cannot account for the increase observed in WT or *pex19*Δ (N_cells_: SD: WT: 245, *hac1*Δ: 270, *pex19*Δ: 263; SD-Ino: WT: 234, *hac1*Δ: 241, *pex19*Δ: 236). b) Induction of lipid stress by growth in the absence of inositol can significantly but only very mildly induce heat shock response as indicated by almost comparable levels of HSE-GFP signal in SD media with or without Inositol (N_cells_: SD: 290, SD-Ino: 171). c-d) Phytosphingosine (PHS) treatment does not increase UPRE-GFP levels (c) or HSE-GFP levels (d) to the same extent as tunicamycin treatment [(c) N_cells_ for WT: DMSO: 466, 0.5µg/ml Tm: 245, 2µg/ml Tm: 267, 20µM PHS: 424; N_cells_ for *hac1*Δ: DMSO: 440, 0.5µg/ml Tm: 262, 2µg/ml Tm: 298, 20µM PHS: 456; (d) N_cells_: DMSO: 272, 0.5µg/ml Tm: 347, 2µg/ml Tm: 402, 20µM PHS: 553].

**Supplemental Figure 3:**
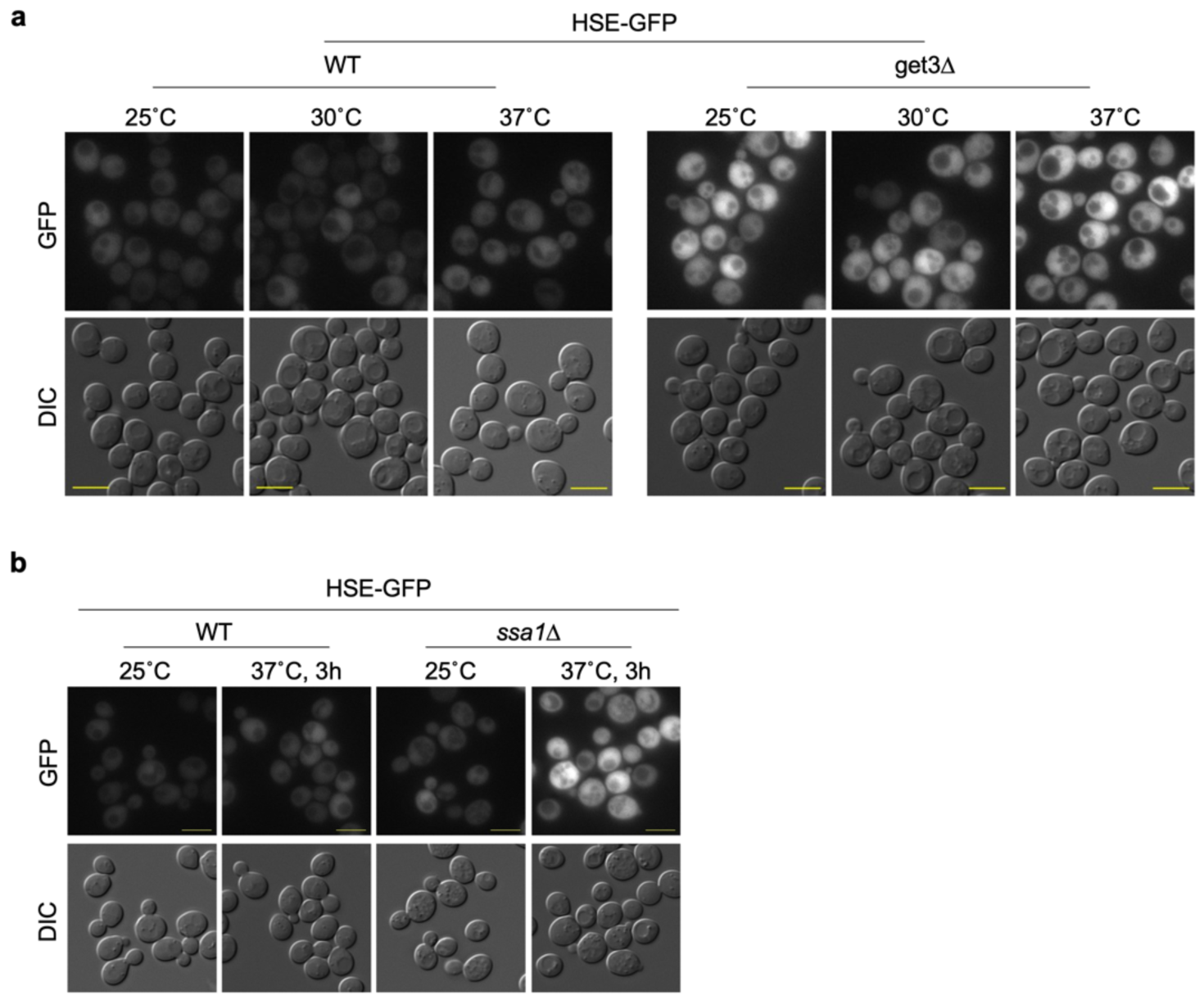
Loss of Get3 or Ssa1 activates heat shock response. a-b) Single Z-slice images showing HSE-GFP signal increases in *get*3Δ and *ssa1*Δ cells in our experimental conditions. Scale bar: 5µm.

**Supplemental Figure 4:**
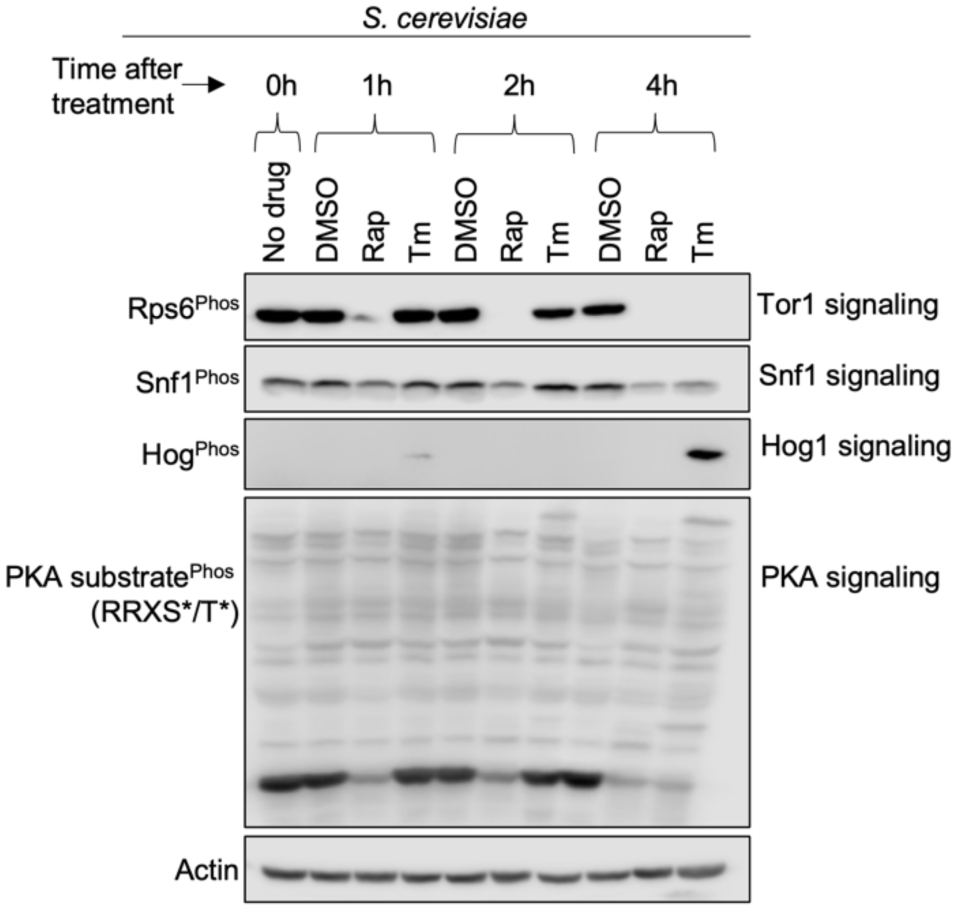
Effects of tunicamycin treatment on multiple signaling pathways. Western blots showing the levels of phosphorylated Rps6, Snf1, and Hog1 to visualize activation of Tor1, Snf1 and Hog1, respectively, at different time points after 0.5µg/mL tunicamycin and 0.5µg/mL rapamycin treatments in comparison to DMSO control. Changes in PKA activity tested by probing for changes in phosphorylation profile of PKA substrate. Actin levels depict equal loading. Overall, the data show that Tor1 inhibition and changes in PKA signaling occur even after 1h of rapamycin treatment, however, these changes are not observed at least until more than 2h of tunicamycin treatment. Hog1 activation is not observed in rapamycin-treated cells, or until more than 2h in tunicamycin-treated samples. Surprisingly Snf1 activation is also not observed after treatment with both drugs under our experimental conditions.

**Supplemental Table 1:**
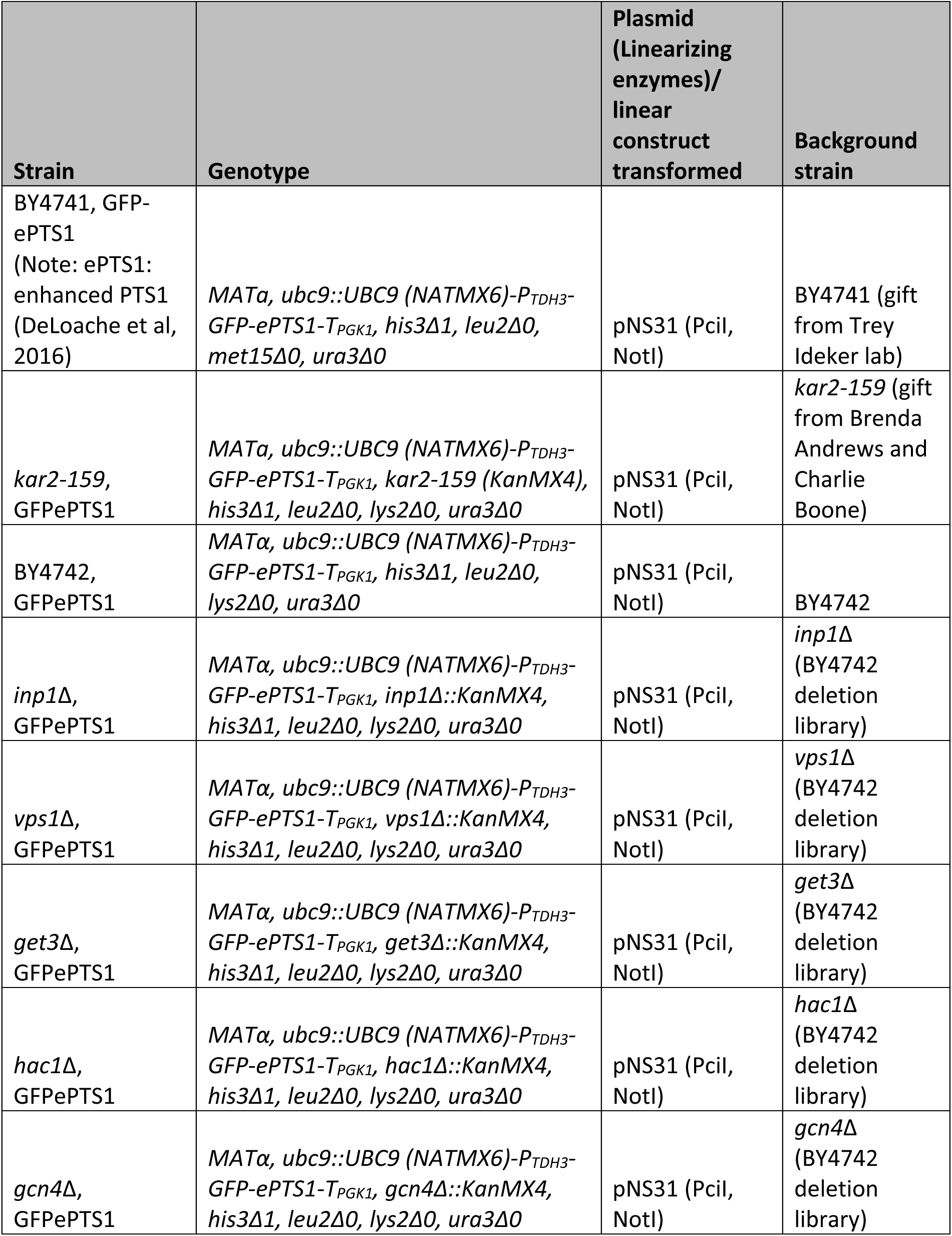
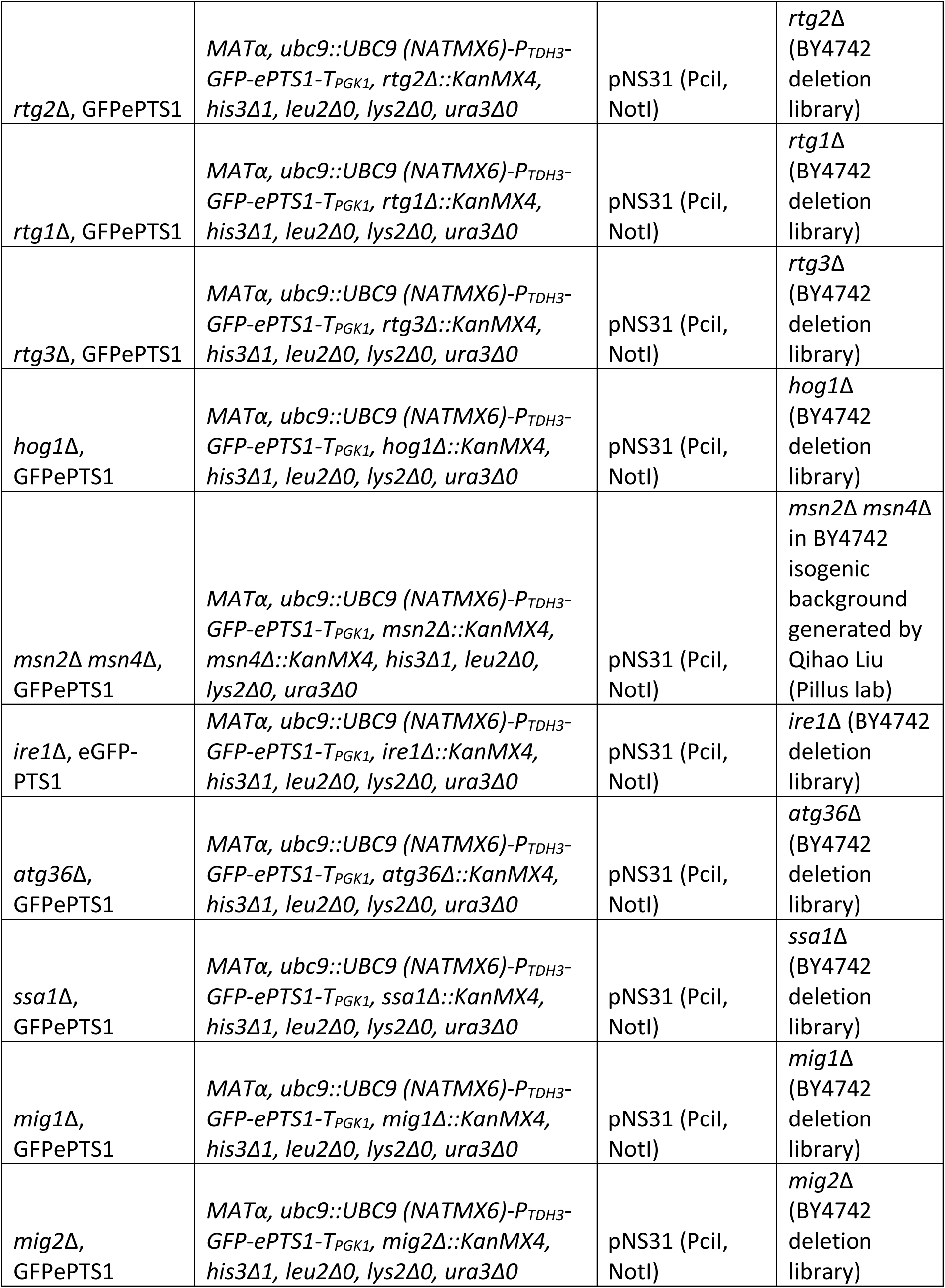
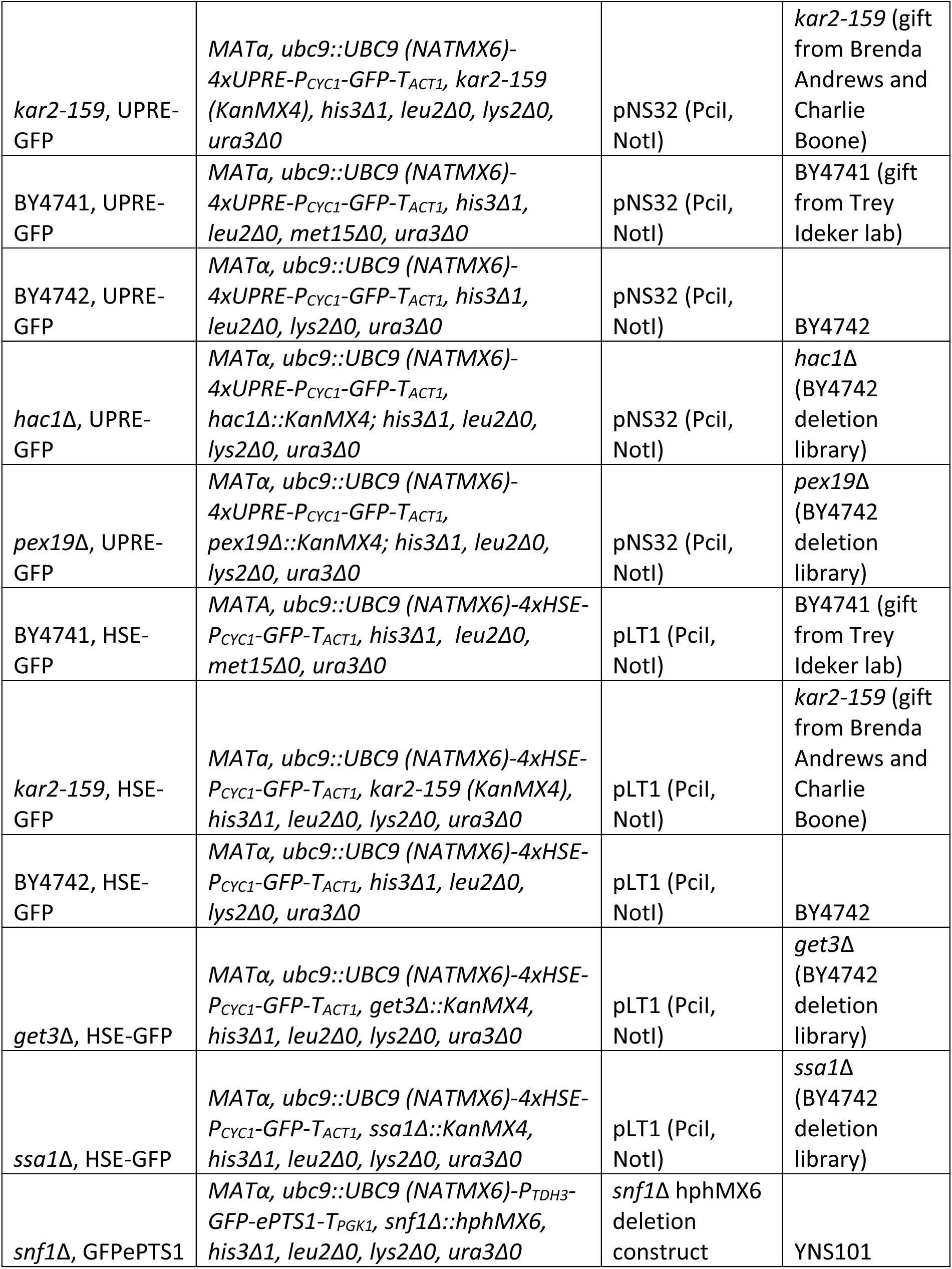
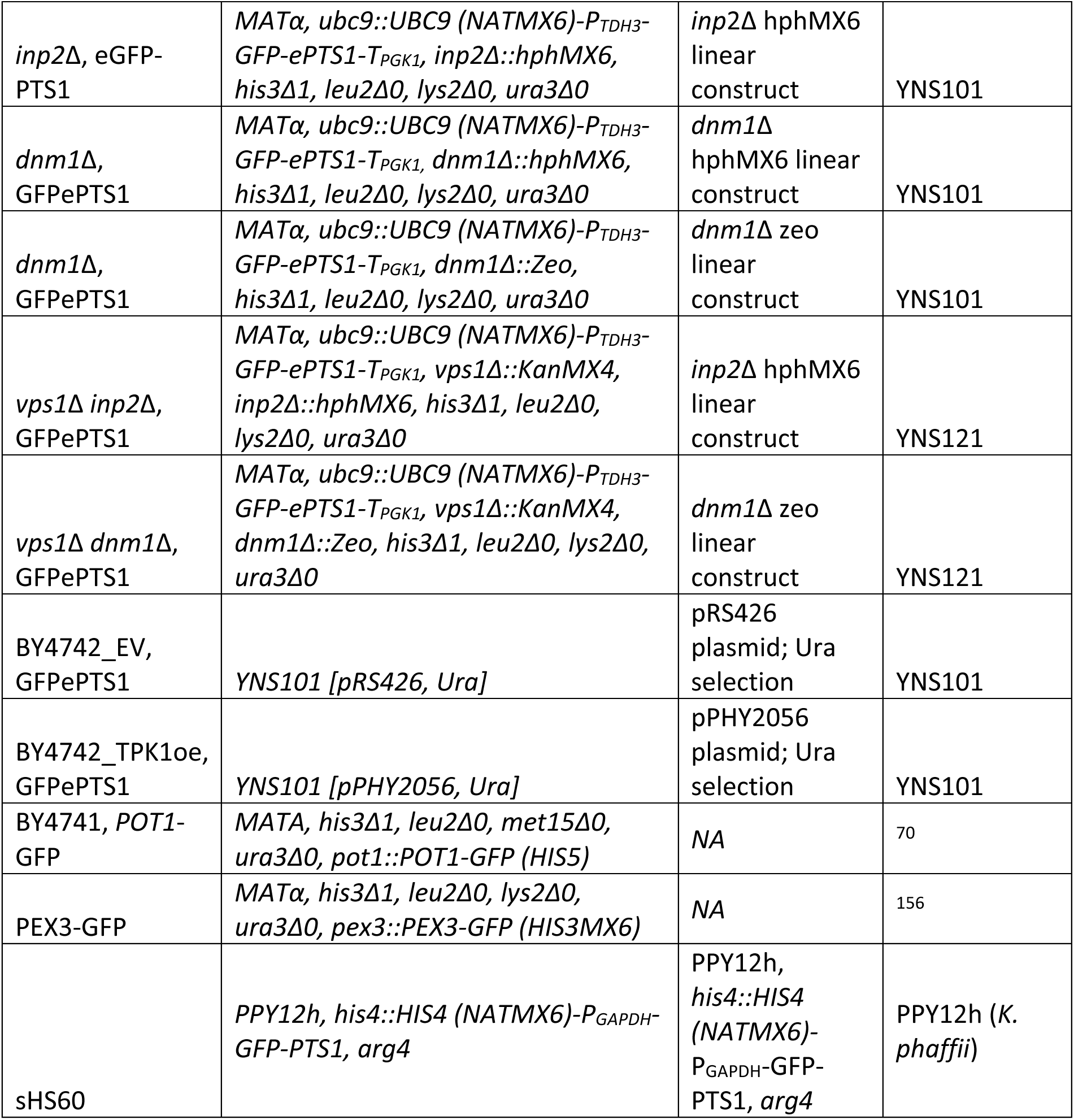
Strain list.

## Abbreviations

CSM: Complete Supplement Mixture
ER: endoplasmic reticulum
DIC: Differential Interference Contrast
GET: Guided entry of tail-anchored proteins
GFP: Green Fluorescent Protein
GSH: glutathione (reduced)
GSSG: oxidized glutathione
HSE: Heat Shock Response Element
*K. phaffii*: *Komagataella phaffii*
PMP: Peroxisomal membrane protein
PO: peroxisome
PTS: peroxisomal targeting signal
*S. cerevisiae*: *Saccharomyces cerevisiae*
SD: synthetic defined
STE: steryl esters
TAG: triacylglycerol
TRX: thioredoxin
ts: temperature sensitive
YE: Yeast extract
YNB: Yeast nitrogen base
UPR: Unfolded Protein Response
UPRE: Unfolded Protein Response Element
WT: Wild type (n)/ wild-type (adj.)

